# Potency, dissociation kinetics and reversibility of fentanyls and nitazenes by naloxone at the μ opioid receptor

**DOI:** 10.1101/2024.06.04.597322

**Authors:** Norah Alhosan, Damiana Cavallo, Marina Santiago, Eamonn Kelly, Graeme Henderson

**Author notes:** **Correspondence:** Graeme Henderson, School of Physiology, Pharmacology and Neuroscience, University of Bristol, Bristol BS8 1TD, UK.

## Abstract

**Background and Purpose:** Fentanyls and nitazenes are μ opioid receptor agonists responsible for a large number of opioid overdose deaths. Here, we compared the potency, dissociation kinetics and antagonism by naloxone at the μ receptor of several fentanyl and nitazene analogues and compared them to morphine and DAMGO.

**Experimental Approach:** *In vitro* assays of G protein activation and signalling and arrestin recruitment were performed. AtT20 cells expressing μ receptors were loaded with a membrane potential dye and changes in fluorescence used to determine agonist potency, dissociation kinetics and susceptibility to antagonism by naloxone. BRET experiments were undertaken in HEK293T cells expressing μ opioid receptors, to assess Gi protein activation and β-arrestin 2 recruitment.

**Key Results:** The rate of agonist dissociation from the μ receptor varied, with morphine, DAMGO, alfentanil and fentanyl dissociating rapidly whereas isotonitazene, etonitazene, ohmefentanyl and carfentanil dissociated slowly. Slowly dissociating agonists were more resistant to antagonism by naloxone. For carfentanil, the slow rate of dissociation was not due to G protein receptor kinase-mediated arrestin recruitment as its rate of dissociation was not affected by inhibition of GRKs with Compound 101. The *in vitro* relative potencies of fentanyls and nitazenes compared to morphine were much lower than that previously observed in *in vivo* experiments.

**Conclusions and Implications:** With fentanyls and nitazenes, that slowly dissociate from the μ opioid receptor, antagonism by naloxone is pseudo competitive. In overdoses involving fentanyls and nitazenes higher doses of naloxone may be required for reversal than those normally used to reverse heroin overdose.

**What is already known:**

- “Fentanyls” and “nitazenes” are potent agonists at the μ opioid receptor.

**What does this study add:**

- Some fentanyls and nitazenes dissociate slowly and are less sensitive to naloxone antagonism

**What is the clinical significance:**

- More naloxone may be required to reverse overdoses involving fentanyls and nitazenes

## 1. Introduction

### 1.1 Fentanyls and nitazenes

For the past 10 years North America has been suffering a synthetic opioid drug epidemic that has resulted in hundreds of thousands of overdose deaths (Centers for Disease Control and Prevention, 2022; Health Infobase, 2024). These opioid overdose deaths primarily involve fentanyls (fentanyl and its analogues). In the UK and mainland Europe nitazenes, rather than fentanyls, are thought to pose a credible threat given the predicted decrease in heroin availability from Afghanistan (Caulkins, Tallaksen *et al*., 2024; Giraudon, Abel-Ollo *et al*., 2024; Holland, Copeland *et al*., 2024;).

### 1.2 Naloxone antagonism

While there are four main types of opioid receptor (μ, δ, κ and NOP) in the nervous system it is through the μ opioid receptor that fentanyls and nitazenes produce their profound physiological effects, including respiratory depression, the main cause of death in overdose. Naloxone, the opioid antidote used to treat overdose, is generally regarded to be a competitive antagonist at the μ opioid receptor (Corbett, Henderson *et al*., 2006). For agonists and antagonists interacting at the same orthosteric binding site on a receptor a competitive antagonist such as naloxone would be expected to reverse all agonists equally given that the antagonist binds to the unbound receptor and prevents agonist binding rather than physically displacing the agonist from the receptor (Ritter, Flower *et al*., 2024). However there have been numerous reports of more naloxone, in the form of multiple or higher doses, being required to reverse overdoses involving fentanyls and nitazenes compared with heroin overdoses (Moe, Godwin *et al*., 2020; Irvine, Oller *et al.,* 2022; Amaducci, Aldy *et al*., 2023). Reduced sensitivity to naloxone has also been observed in animal studies using sub lethal doses of fentanyls (Hill, Santhakumar *et al*., 2020; Elder, Varshneya *et al*., 2023)

### 1.3 Aims

In this study, we compared the potency and dissociation kinetics of a range of fentanyls (alfentanil, fentanyl, sufentanil, ohmefentanyl and carfentanil) and nitazenes (isotonitazene, etonitazene) at the μ opioid receptor *in vitro* and compared them to the prototypic opioid agonists morphine and D-Ala^2^, *N*-MePhe^4^, Gly-ol]-enkephalin (DAMGO). We also examined the ability of naloxone to antagonise the opioid agonists. Our results indicated that some fentanyls and nitazenes were less susceptible to naloxone antagonism than morphine and that this correlated with how slowly those agonists dissociated from the receptor.

## 2. Methods

### 2.1 Cell Culture

‘Empty’ Flp-In modified AtT20 cells (AtT20FlpInWT) (ATCC CRL-1795; RRID:CVCL_4109) and Flp-In modified AtT20 cells recombinantly expressing the human μ opioid receptor (AtT20FlpInMOR) were obtained from Prof M Connor, Macquarie University, Australia. Cells were maintained in 75 cm flasks at 37°C and 5% CO_2_ in Dulbecco’s Modified Eagle’s Medium (DMEM) containing L-Glutamine supplemented with 10% foetal bovine serum (FBS), 50 U mL^−1^ penicillin, 0.5 mg mL^−1^ streptomycin and 80 μg mL^−1^ hygromycin B. Human embryonic kidney 293T (HEK293T) cells were grown in 10 cm dishes in DMEM supplemented with 10% FBS and 50 U mL^−1^ penicillin and 0.5 mg mL^−1^ streptomycin and incubated in 5% CO_2_ at 37°C.

### 2.2 Membrane potential assay

The protocol used was a minor modification of that previously described (Knapman and Connor, 2015). When μ opioid receptor-expressing AtT20 cells reached ≈ 90% confluency, they were detached by trypsinization, resuspended in Leibovitz’s L-15 medium and plated into black, clear flat bottom 96 well assay plates coated with 0.01% poly-L-lysine solution. Each well received 90 μL of cell suspension and the plate incubated overnight in air at 37⁰C. One hour prior to the experiment, cells were loaded with 90 μL of fluorescent blue membrane potential dye (supplied in the FLIPR Membrane Potential Assay Kit, Molecular Devices). Membrane potential dye and drug dilutions were prepared in a low potassium buffer (NaCl 145mM, HEPES 22 mM, Na2HPO_4_ 0.338 mM, NaHCO_3_ 4.17 mM, KH2PO4 0.441 mM, MgSO4 0.407 mM, MgCl_2_ 0.493 mM, CaCl_2_ 1.26 mM, Glucose 5.56 mM, pH 7.4). Use of low potassium buffer reduced the extracellular potassium concentration in the solution bathing the cells during the experimental recordings from 5.6 mM to 2.9 mM, making the potassium equilibrium potential more negative, thus enhancing the amplitude of the opioid agonist-induced hyperpolarising responses. Fluorescence was measured from cells maintained at 37°C using a FlexStation 3 Multi-Mode Microplate Reader (Molecular Devices). Cells were excited at a wavelength of 530 nm and emission measured at 565 nm, with readings taken every 2 s. In control experiments there was no decrease in the fluorescence measured in this way over a 500 s experimental run as might have occurred with dye bleaching. Background fluorescence in wells with cells only or dye only was low and regarded as negligible.

Drug additions were performed using the robotic function of the FlexStation 3. All drugs were added in a volume of 10 µL and ejected at a speed of 16 µL/sec. The tip of the drug-containing pipette was placed at a height equivalent to the upper surface of the bathing fluid (180 μL for the first addition and 190 μl for the second). Following drug addition the fluid in the well was ‘stirred’ by removing and re-injecting 10µL of the bathing fluid three times as per the trituration setting of the FlexStation. In preliminary experiments examining the rate of potassium channel block by barium (a response that should have no intrinsic lag time) we estimated that the minimum response time measurable by the FlexStation following drug addition and mixing was 5.8 ± 0.4s

In the membrane potential assay experiments we first examined the amplitude of the fluorescence signal from each well to ensure consistency of cell density, the condition of the cells and dye loading within and between experiments. All experiments were performed on wells that exhibited an initial fluorescence signal between 500 – 900 RFU. To control for differences in the basal level of the fluorescent signal between wells the amplitude of subsequent drug-evoked responses was calculated as the percentage change from baseline, pre-drug addition, fluorescence readings in each well. Responses from wells that were injected with buffer alone rather than drug were subtracted to compensate for the small injection artefacts observed in some experiments. The change in the signal produced by the addition of buffer alone was ≤5% of the baseline.

### 2.3 Bioluminescence Resonance Energy Transfer assays

To determine the relative ability of the opioid agonists to activate Gα_i_ G proteins and β-arrestin 2 translocation to the receptor, BRET^2^-based assays were used as described previously (Hill, Disney *et al*., 2018; Ramos-Gonzalez, Groom *et al*., 2023). For Gi activation the assay monitored the separation of Gα_i1_ and Gγ_2_ whereas for arrestin translocation the assay measured μ opioid receptor and β-arrestin 2 association.

HEK 293T cells were transiently transfected with the appropriate constructs (for Gi activation - rat HA-μ opioid receptor, Gα_i1_-*Renilla* luciferase II (RlucII) and GFP10-Gγ_2_; and for arrestin translocation - human μ opioid receptor-RlucII and β-arrestin-2-GFP). They were then incubated for a further 48 h before the BRET assays were conducted. Immediately prior to each assay, cells were resuspended in clear DMEM and then transferred to a 96-well plate at 90 μL per well. Measurements of BRET were made at 37°C. Coelenterazine 400a, at a final concentration of 5 μM, was injected 5 s prior to reading the cell plate. BRET measurements were made on a CLARIOstar Omega plate reader (BMG LABTECH, Ortenberg, Germany) using 515 ± 30 nm (acceptor) and 410 ± 80 nm (donor) filters. BRET signals were determined as the ratio of the light emitted by acceptors (GFP10) over donor (RlucII). For Gi activation BRET measurements were taken 2 min after agonist application and for β-arrestin 2 association 10 min after agonist application. Agonist application resulted in a rapid decrease in the BRET signal between Gα_i1_-RlucII and GFP10-Gγ_2_ and an increase in the BRET signal between μ opioid receptor-Rluc and β-arrestin-2-GFP. The ratio of the signal from the acceptor and donor was then calculated. Data were expressed either as percentage decrease for the Gi activation assay, or as raw data with basal subtracted for the β-arrestin 2 recruitment assay.

### 2.4 Bias Calculations

IUPHAR guidelines for estimating GPCR ligand bias were followed to calculate bias (Kolb, Kenakin *et al*., 2022). Bias was calculated using two methods: using the operational model (ΔΔLog t/K_A_) and the Log (E_max_/EC_50_) model. Concentration-response curves for each drug were generated in the G protein activation and β-arrestin 2 recruitment assays and Log (E_max_/EC_50_) values were generated using 3-parameter nonlinear regression fit. In each assay the mean Log (E_max_/EC_50_) of each agonist was compared to the reference full agonist DAMGO to obtain Δ Log (E_max_/EC_50_). Then for each agonist the Δ Log (E_max_/EC_50_) values between G protein activation and β-arrestin 2 recruitment assays were compared to acquire the ΔΔ Log (E_max_/EC_50_). For Log(τ/K_A_) analysis, agonist concentration-response curves for G protein activation and β-arrestin 2 recruitment were fitted to the Black-Leff operational model (Black & Leff, 1983) to generate Log(τ/K_A_) (Transduction Ratio) values (Kolb et al., 2022). A similar calculation to the one above for ΔΔ Log (E_max_/EC_50_) was undertaken to generate ΔΔLog (t/K_A_).

### 2.6 Experimental design and data analysis

This manuscript complies with BJP’s recommendations and requirements on experimental design and analysis (Curtis, Alexander *et al*., 2018; Curtis, Alexander *et al*., 2022).

In the membrane potential and BRET assays the order of addition of different drugs and their concentrations were randomised within individual experiments and across a series of experiments. Accordingly, the location of the basal or vehicle controls was different for each experiment. Blinding was not undertaken due to the complexity of the plate assays with multiple agonists and concentrations. Drug addition and data accumulation were automated by the software controlling the FlexStation and CLARIOstar 96 well plate readers. Raw data were downloaded into Excel spreadsheets for subsequent analysis. Membrane potential and BRET assays were conducted in duplicates. The mean of each duplicate was calculated and considered as n = 1. A priori sample size estimation indicated that a sample size of <5 would provide sufficient power for both membrane potential and BRET experiments. We therefore used a sample size of n = 5 in all experiments.

Data analysis, including statistical testing, was carried out in GraphPad Prism 9.0. For all assays, data points were excluded where their value was >3 x S.D. different from the mean of the other values. Concentration-response and naloxone reversal experiments with fentanyls and nitazenes were performed in two sets at different times. To allow comparison between each we included morphine as a control in both. To construct concentration-response curves for each agonist, responses were normalised to that produced by morphine (1 μM) in the same set of experiments to remove any potential change in maximum response amplitude between the two sets of experiments.

pEC_50_, pIC_50_ and maximum response values were obtained by fitting concentration-response curves from each experiment individually by non-linear regression (the initial value for no drug was constrained to zero or 100% as appropriate) and then combining values to give mean values ± S.E.M (n=5). In the analysis of the concentration-response curves from the BRET experiments it was not possible to fit the data for morphine, a weak partial agonist in the b-arrestin 2 translocation assay, without constraining the Hill slope to 1 and so for consistency the Hill slope was constrained to 1 for all agonists.

Apparent dissociation half-times were obtained by fitting the decay of the agonist response after naloxone (10μM) addition in each individual experiment to a custom equation downloadable to GraphPad from Pharmechanics. This took into account a variable ‘delay’ period following naloxone addition that was not different from the pre-drug baseline followed by the exponential decay to steady state. In each experiment the goodness of fit was r ≥ 0.8. Note: with regards to the rate of agonist dissociation from the receptors (see Figure 4 and Table 1) we use the term ‘apparent’ half time of agonist dissociation to indicate that although a high concentration of the antagonist naloxone has been added the agonist has not been washed out and so we cannot exclude that a small amount of agonist rebinding to the receptors may occur thus slowing the rate of reversal.

**Table 1.**
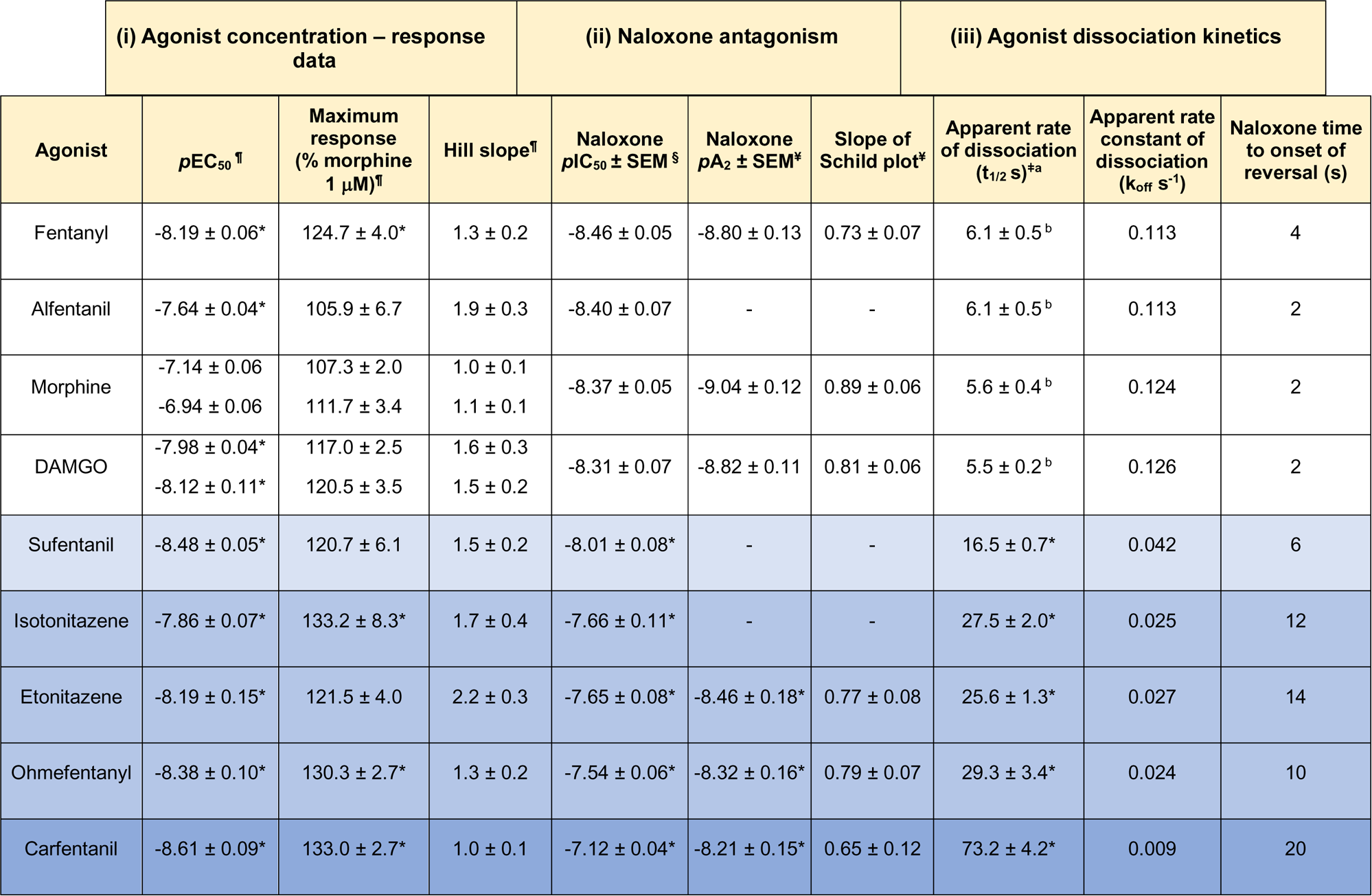
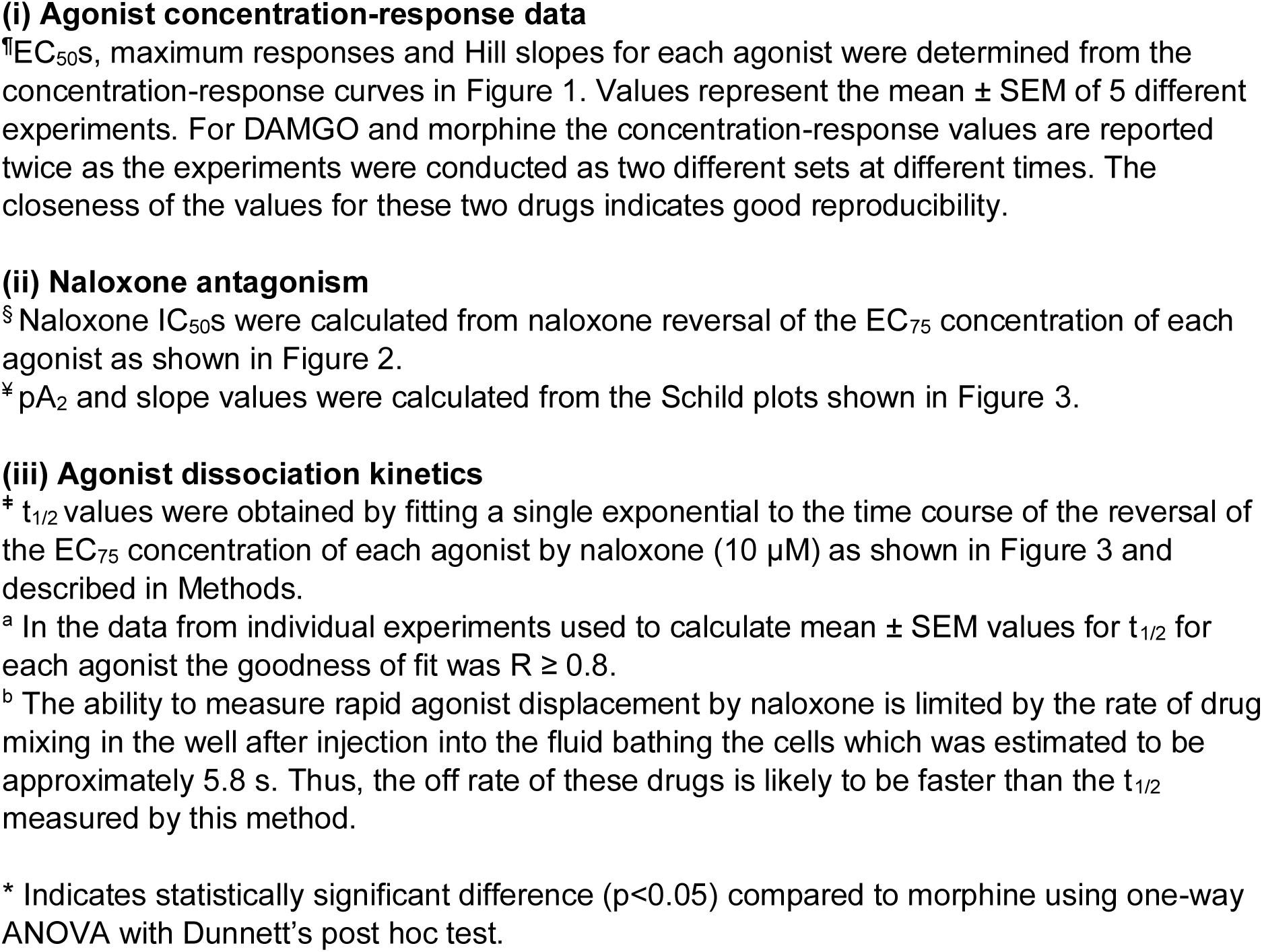
Data obtained from agonist concentration-response, naloxone reversal and agonist dissociation kinetic experiments.

| (i) Agonist concentration – response data |  |  |  | (ii) Naloxone antagonism |  |  | (iii) Agonist dissociation kinetics |  |  |
| --- | --- | --- | --- | --- | --- | --- | --- | --- | --- |
| Agonist | $pEC_{50}^{\dagger}$ | Maximum response (% morphine 1 μM)<sup>†</sup> | Hill slope<sup>†</sup> | Naloxone $pIC_{50} \pm SEM^{\S}$ | Naloxone $pA_2 \pm SEM^{\P}$ | Slope of Schild plot<sup>‡</sup> | Apparent rate of dissociation ($t_{1/2}$ s)<sup>*a</sup> | Apparent rate constant of dissociation ($k_{off}$ s<sup>-1</sup>) | Naloxone time to onset of reversal (s) |
| Fentanyl | $-8.19 \pm 0.06^*$ | $124.7 \pm 4.0^*$ | $1.3 \pm 0.2$ | $-8.46 \pm 0.05$ | $-8.80 \pm 0.13$ | $0.73 \pm 0.07$ | $6.1 \pm 0.5^b$ | 0.113 | 4 |
| Alfentanil | $-7.64 \pm 0.04^*$ | $105.9 \pm 6.7$ | $1.9 \pm 0.3$ | $-8.40 \pm 0.07$ | - | - | $6.1 \pm 0.5^b$ | 0.113 | 2 |
| Morphine | $-7.14 \pm 0.06$<br>$-6.94 \pm 0.06$ | $107.3 \pm 2.0$<br>$111.7 \pm 3.4$ | $1.0 \pm 0.1$<br>$1.1 \pm 0.1$ | $-8.37 \pm 0.05$ | $-9.04 \pm 0.12$ | $0.89 \pm 0.06$ | $5.6 \pm 0.4^b$ | 0.124 | 2 |
| DAMGO | $-7.98 \pm 0.04^*$<br>$-8.12 \pm 0.11^*$ | $117.0 \pm 2.5$<br>$120.5 \pm 3.5$ | $1.6 \pm 0.3$<br>$1.5 \pm 0.2$ | $-8.31 \pm 0.07$ | $-8.82 \pm 0.11$ | $0.81 \pm 0.06$ | $5.5 \pm 0.2^b$ | 0.126 | 2 |
| Sufentanil | $-8.48 \pm 0.05^*$ | $120.7 \pm 6.1$ | $1.5 \pm 0.2$ | $-8.01 \pm 0.08^*$ | - | - | $16.5 \pm 0.7^*$ | 0.042 | 6 |
| Isotonitazene | $-7.86 \pm 0.07^*$ | $133.2 \pm 8.3^*$ | $1.7 \pm 0.4$ | $-7.66 \pm 0.11^*$ | - | - | $27.5 \pm 2.0^*$ | 0.025 | 12 |
| Etonitazene | $-8.19 \pm 0.15^*$ | $121.5 \pm 4.0$ | $2.2 \pm 0.3$ | $-7.65 \pm 0.08^*$ | $-8.46 \pm 0.18^*$ | $0.77 \pm 0.08$ | $25.6 \pm 1.3^*$ | 0.027 | 14 |
| Ohmefentanyl | $-8.38 \pm 0.10^*$ | $130.3 \pm 2.7^*$ | $1.3 \pm 0.2$ | $-7.54 \pm 0.06^*$ | $-8.32 \pm 0.16^*$ | $0.79 \pm 0.07$ | $29.3 \pm 3.4^*$ | 0.024 | 10 |
| Carfentanil | $-8.61 \pm 0.09^*$ | $133.0 \pm 2.7^*$ | $1.0 \pm 0.1$ | $-7.12 \pm 0.04^*$ | $-8.21 \pm 0.15^*$ | $0.65 \pm 0.12$ | $73.2 \pm 4.2^*$ | 0.009 | 20 |

All data were tested for normality using the D’Agostino & Pearson test and graphically using Q-Q plot and bar graphs. To test for statistical differences between pEC_50_, E_max_, pIC_50_ and dissociation half-time values, all ligands were compared to morphine (membrane potential assay) or DAMGO (BRET assay) as the reference agonist using one-way ANOVA. Post-hoc Dunnett’s multiple comparisons were conducted only if the results showed a statistical significance (F value achieved a P value of < 0.05) and no inhomogeneity in variance. For the correlation analyses nonparametric Spearman test was used as the data were not normally distributed and a two-tailed P value of <0.05 taken to indicate significance.

### 2.7 Materials

The drugs used were alfentanil hydrochloride (Cayman Chemicals), carfentanil and ohmefentanyl (Toronto Research Chemicals), Compound 101 (Hello Bio), [D-Ala^2^, N-MePhe^4^, gly-ol] enkephalin acetate (DAMGO) (Bachem and Sigma-Aldrich), etonitazene hydrochloride fentanyl citrate, naloxone hydrochloride, U69593 (Sigma-Aldrich), isotonitazene (Cayman Chemicals), morphine hydrochloride (Macfarlan Smith), nociceptin (Biotechne) and SNC80 (Tocris). Drugs were made up as stocks in deionised water or DMSO and subsequently diluted in the appropriate experimental buffer.

### 2.8 Nomenclature of targets and ligands

Key protein targets and ligands in this article are hyperlinked to corresponding entries in http://www.guidetopharmacology.org, and are permanently archived in the Concise Guide to PHARMACOLOGY 2021/22 (Alexander, Christopoulos *et al*., 2021).

## 3. Results

### 3.1 Opioid agonist concentration-response relationships in the AtT20 cell membrane potential assay

The opioid agonists fentanyl, alfentanil, sufentanil, ohmefentanyl, carfentanil, etonitazene and isotonitazene, as well as the prototypic μ opioid receptor agonists DAMGO and morphine, each produced a concentration-dependent decrease in fluorescence in μ opioid receptor-expressing AtT20 cells loaded with membrane potential dye indicative of membrane hyperpolarisation following G protein-mediated GIRK channel activation (Figure 1). The rank order of potency was carfentanil ≥ sufentanil ≥ ohmefentanyl > fentanyl = etonitazene ≥ DAMGO > isotonitazene ≥ alfentanil > morphine (see Table 1 for pEC_50_ values). In this assay the relative potencies of the fentanyls and nitazenes compared to morphine were lower than might be expected from *in vivo* antinociception experiments (Suzuki & El Haddad, 2017; Hasegawa *et al*., 2022). Fentanyl, isotonitazene, ohmefentanyl, and carfentanil evoked a higher maximum response than morphine (Figure 1g & h and Table 1).

**Figure 1.**
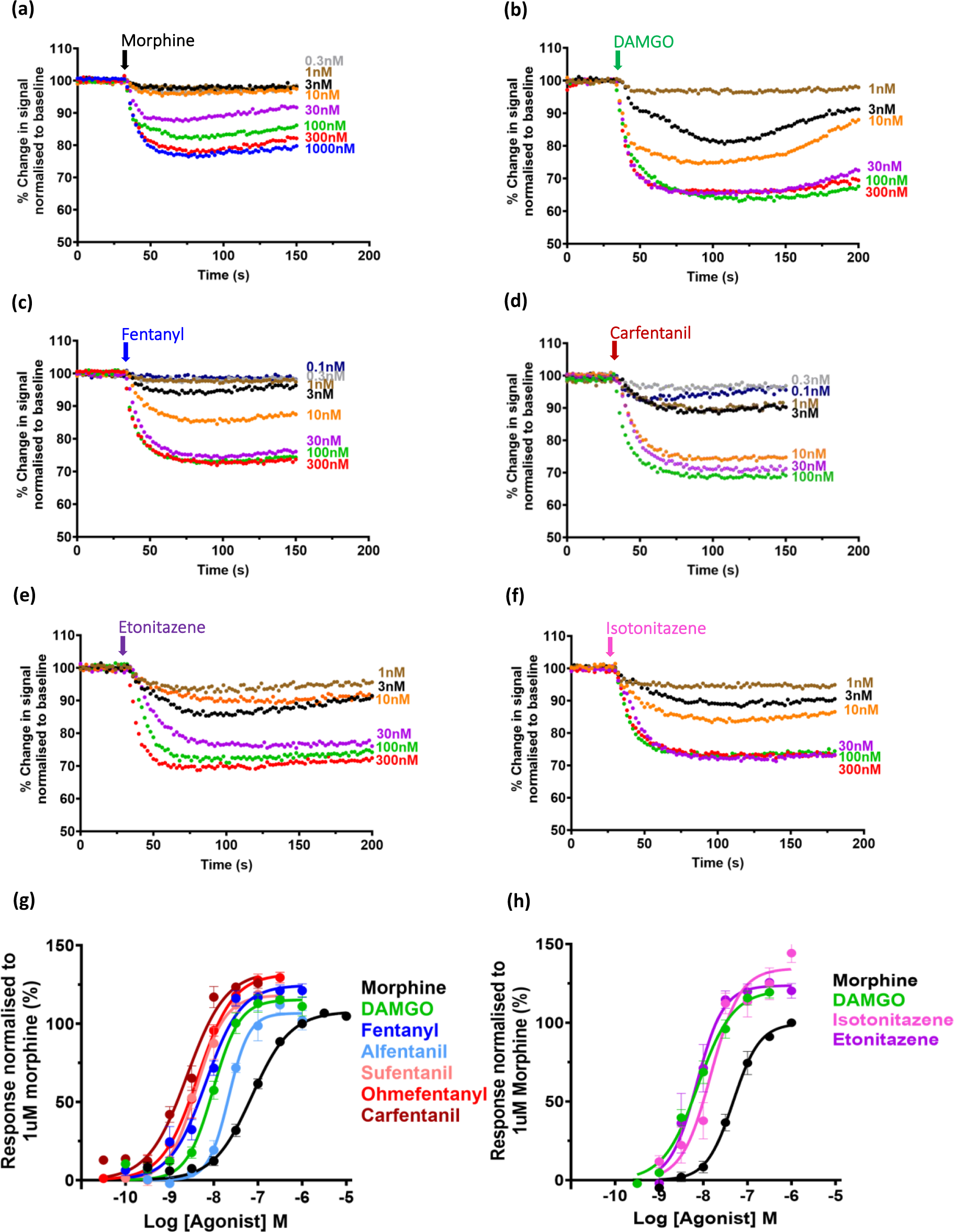
Concentration-response relationships for opioid-induced hyperpolarisation of μ opioid receptor-expressing AtT20 cells. (**a** – **f**) Typical experimental traces for the change in membrane potential dye fluorescence signal produced by increasing concentrations of six opioid agonists - morphine, DAMGO, fentanyl, carfentanil, etonitazene and isotonitazene. Fluorescence values have been normalised to the pre-drug baseline. For each agonist the data are from the same experiment and are typical of responses obtained in five experiments for each agonist. (**g** & **h**) Log concentration-response curves for the decrease in fluorescence induced by all the opioid agonists tested. Data for DAMGO, fentanyl, alfentanil, sufentanil, ohmefentanyl and carfentanil were obtained in a different set of experiments from those for etonitazene and isotonitazene. To facilitate comparison of all opioid agonists, morphine and DAMGO were included in both sets of experiments and changes in fluorescence normalised to the response induced by 1 μM morphine. Data shown are means ± SEM, *n* = 5 for each drug. Log concentration-response curves were constructed using nonlinear regression with the bottom of the curve constrained to zero. Calculated values of agonist pEC_50_, maximum response relative to morphine, Hill slope and relative potency to morphine are given in Tables 1 and 3.

### 3.2 Antagonism of opioid agonists by naloxone

#### 3.2.1 Reversal

In the emergency treatment of human opioid overdose the antagonist naloxone is administered after the response to the agonist has developed. Therefore we first sought to mimic this situation in our *in vitro* experiments by administering each opioid agonist, allowing the response to reach steady state, and then administering naloxone (see Figure 2a-f). Each agonist was added at its EC_75_ concentration and the ability of a range of naloxone concentrations to reverse the agonist response was determined. The order of potency of naloxone to reverse the opioid agonists was morphine = DAMGO = fentanyl = alfentanil > sufentanil > etonitazene = isotonitazene > ohmefentanyl >> carfentanil (Figure 2g & h; see Table 1 for pIC_50_ values for naloxone).

**Figure 2.**
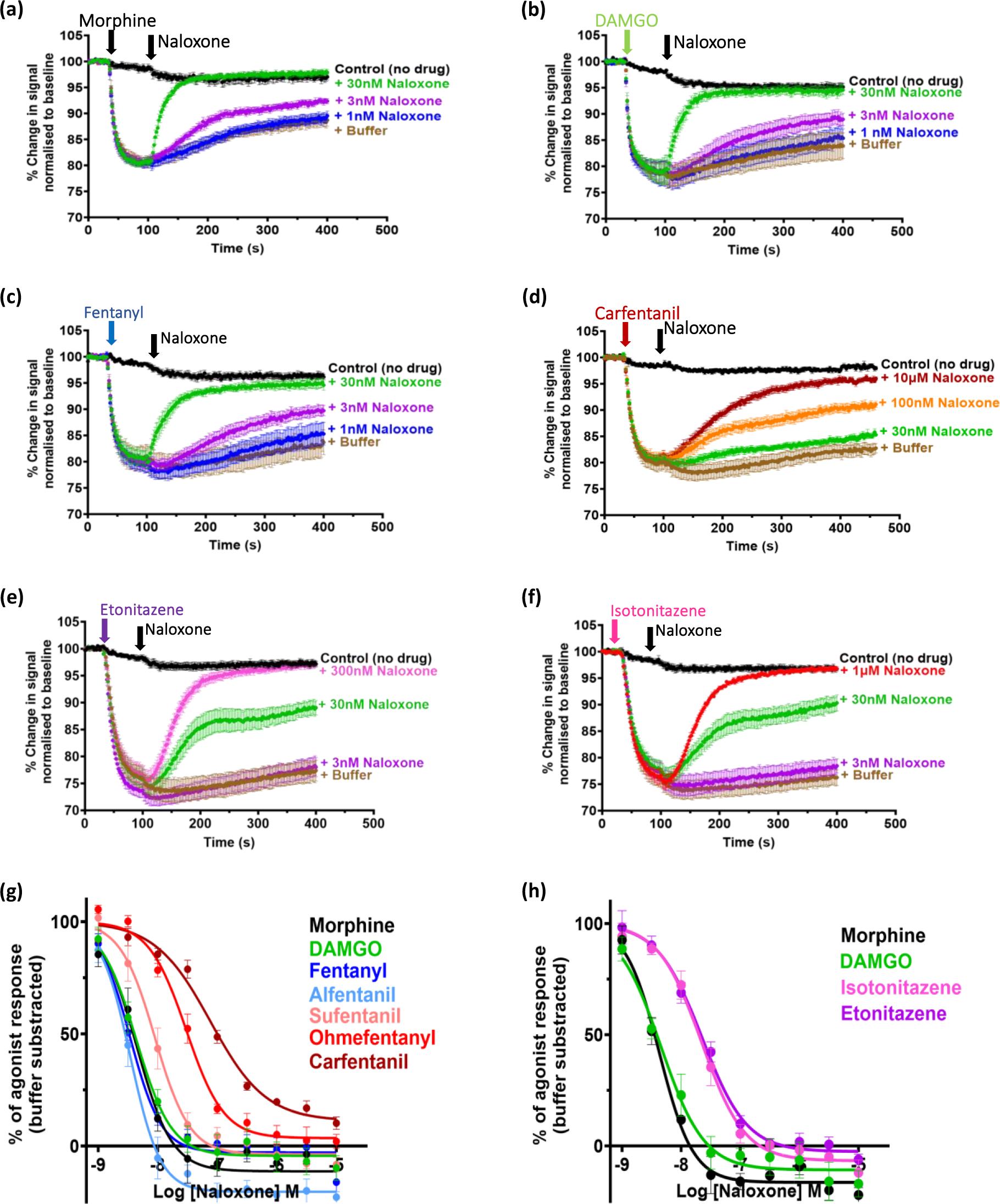
Concentration-response relationships for naloxone reversal of opioid-induced hyperpolarisation of μ opioid receptor-expressing AtT20 cells. (**a** – **f**) Pooled experimental data showing the change in membrane potential dye fluorescence signal produced by the EC_75_ concentration of each of six opioid agonists - morphine, DAMGO, fentanyl, carfentanil, etonitazene and isotonitazene – and the subsequent reversal by addition of three concentrations of naloxone. Traces represent the mean ± SEM of 5 individual experiments for each drug. The black line ‘Control (no drug)’ represents no agonist and no naloxone addition, only equivalent volume of vehicle injections. (**g** & **h**) Log concentration-response curves for the reversal by naloxone of each opioid agonist obtained in experiments similar to those shown in (**a** – **d)**. Data for DAMGO, fentanyl, alfentanil, sufentanil, ohmefentanyl and carfentanil shown in (**g)** were obtained in a different set of experiments from those for etonitazene and isotonitazene shown in (**h)**. To facilitate comparison of all opioid agonists tested, morphine and DAMGO were included in both sets of experiments Data shown are means ± SEM, *n* = 5 for each drug. Log concentration-response curves were fitted using nonlinear regression with the top constrained to 100%. Calculated values of naloxone pIC_50_ against each agonist are given in Table 1.

#### 3.2.2 Competitive antagonism

Estimation of the pA_2_ and thus the equilibrium dissociation constant (KD) of a competitive antagonist requires that binding of the antagonist to the receptor is at equilibrium before addition of the agonist to compete with the antagonist for binding. In theory, for competitive antagonism at the same receptor the antagonist pA_2_ should be independent of the agonist as the antagonist binds to the receptor when it is unoccupied by the agonist (Kenakin, 1982). We therefore exposed μ opioid receptor-expressing AtT20 cells to increasing concentrations of naloxone for 30 min at 37°C prior to determining the log concentration-response relationship of each agonist and used these data to measure the concentration ratio of the agonist (Figure 3). Naloxone (3 – 300 nM) produced increasing parallel shifts to the right of the concentration-response curve of each agonist with no decrease in maximum response. However the degree of rightward shift produced by naloxone was not the same for each agonist (Figure 3a-f) and subsequent Schild analysis revealed that the naloxone pA_2_ was not the same for each agonist (Figure 3g and Table 1). For DAMGO and morphine, and fentanyl, the pA_2_ values for naloxone were in the range −9.04 to −8.80 (KD 1.0 to 1.58 nM) which is similar to the Ki value of 1.3 nM reported from radioligand binding studies on human µ opioid receptors (Toll, Berzetei-Gurske *et al*., 1998). However, for the other agonists the naloxone pA_2_ was greater indicating that they were less sensitive to naloxone antagonism. The rank order of naloxone sensitivity was DAMGO = morphine = fentanyl > etonitazene > ohmefentanyl > carfentanil. While the slopes of the Schild plots for DAMGO and morphine were 0.8 and 0.9, respectively, they were statistically different from unity as determined by the extra-sum-of-squares F test. For the other agonists, the slope was even lower, with carfentanil displaying the lowest slope of 0.65 (Table 1). This may be indicative of the interaction between naloxone and the fentanyl and nitazene agonists not being truly competitive in nature (see Discussion) or of the agonists and naloxone acting on more than one subtype of opioid receptor. Therefore we have not converted the observed pA_2_ values to KD values for the fentanyl and nitazene analogues (Kenakin 1982).

**Figure 3.**
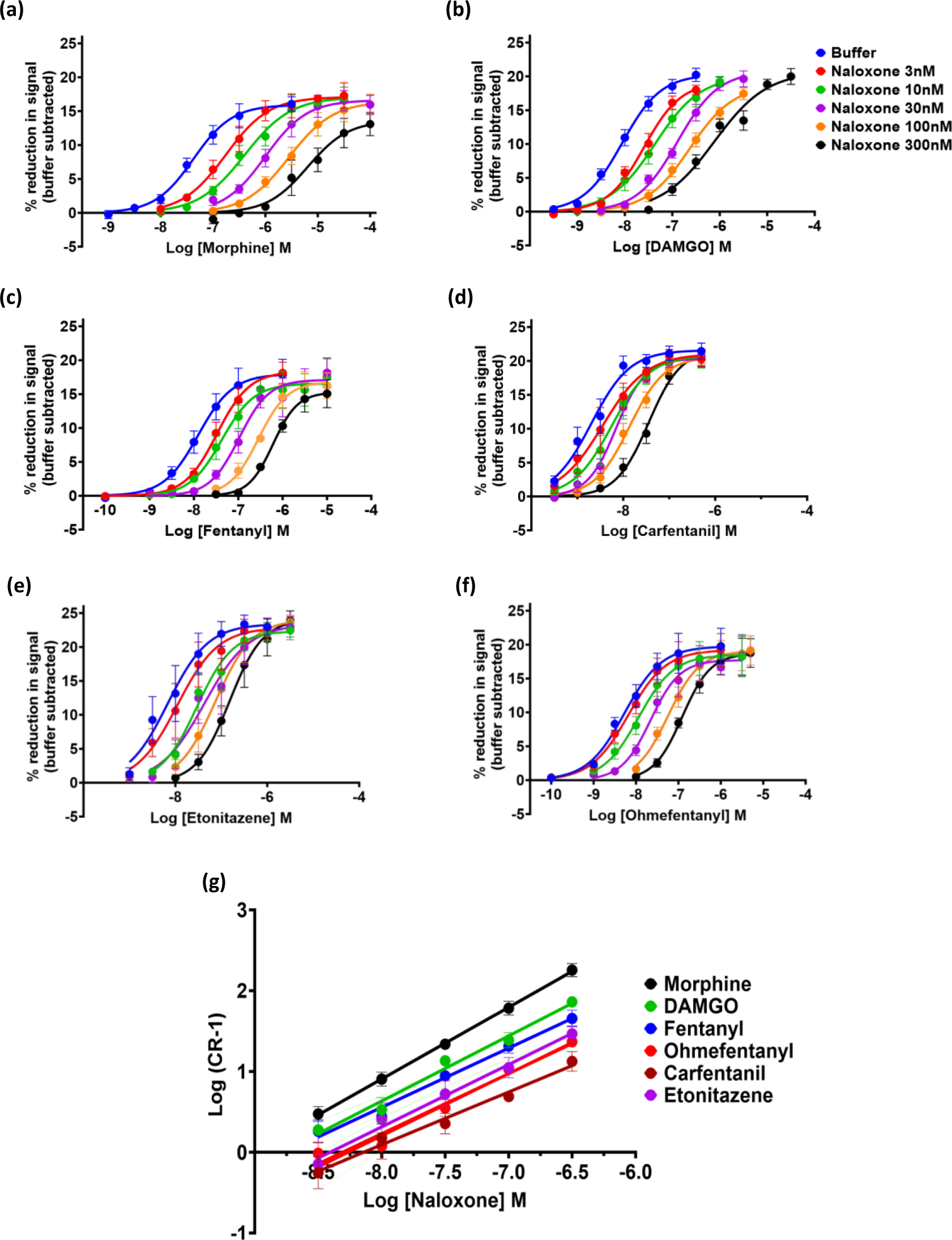
Schild analysis of the antagonism by naloxone of opioid-induced hyperpolarisation of μ opioid receptor-expressing AtT20 cells. For the Schild analysis cells were exposed to naloxone at various concentrations for 30 mins prior to addition of a range of concentrations of each opioid agonist. (**a** – **f**)) Log concentration-response curves for six opioid agonists - morphine, DAMGO, fentanyl, ohmefentanyl, carfentanil and etonitazene – in the absence and presence of naloxone. Each data point represents the mean +/− SEM of n = 5 observations. Log concentration-response curves were fitted using nonlinear regression with the bottom constrained to zero. (**g**) Schild plots of log (concentration ratio – 1) against log naloxone concentration for each of the opioid agonists tested. Data shown are means ± SEM, *n* = 5 for each drug. The data were fitted using simple linear regression. Calculated values of naloxone pA_2_ are given in Table 1.

To exclude the possibility that the opioid agonists were activating more than one type of opioid receptor in the AtT20 cells we examined whether δ, κ or NOP opioid receptors were endogenously expressed in the parent cell line, AtT20FlpInWT, in which the μ opioid receptor had subsequently been recombinantly expressed. A commercial transcriptome analysis performed by Macrogen^©^ had previously indicated that the AtT20FlpInWT cells did not endogenously express δ or κ opioid receptors and only endogenously expressed NOP opioid receptors at a very low level (the full transcriptome analysis data for AtT20FlpInWT cells have been made publicly available by Prof M. Connor, Macquarie University, Australia at https://doi.org/10.25949/21529404.v1). We attempted to confirm the absence of these opioid receptors by examining whether AtT20FlpInWT cells responded to opioid agonists selective for δ, κ or NOP opioid receptors. When we exposed AtT20FlpInWT cells (i.e. the parent cell line not recombinantly expressing the μ opioid receptor) to SNC80 (1 μM), U69593 (10 μM) or nociception (1 μM), agonists at δ, κ and NOP opioid receptors respectively, there was no decrease in fluorescence in cells loaded with membrane potential dye. This is consistent with these cells not endogenously expressing significant amounts of δ, κ or NOP opioid receptors that couple to GIRK channels. In addition, we sought to exclude the possibility that the high concentrations of carfentanil or etonitazene used to overcome antagonism by naloxone (Figure 3), might have off target effects at non opioid receptors in AtT20 cells so reducing the effectiveness of naloxone. When we exposed AtT20FlpInWT cells to carfentanil (100 nM) or etonitazene (300 nM) there was no change in fluorescence in cells loaded with membrane potential dye indicating that at even at the high concentrations carfentanil and etonitazene were devoid of off target actions that affected membrane potential.

### 3.3 Off rate of agonist binding

To examine whether the reduced sensitivity to reversal by naloxone of some fentanyls and nitazenes (see 3.2.1 above) might be due to slow agonist dissociation from the receptor we measured the apparent rate of agonist dissociation by allowing the response to the EC_75_ concentration of each agonist to reach steady state before applying a receptor supersaturating concentration of naloxone (10 μM)(Figure 4). The decay phase of the agonist response in the presence of naloxone was fitted to a single exponential and the apparent t_1/2_ of dissociation determined. For fentanyl, alfentanil, morphine, and DAMGO, the apparent rate of agonist dissociation was similar to the response time of our assay procedure, as the values of apparent t_1/2_ of dissociation obtained were similar to the estimated equilibrium time for mixing following drug injection into the medium bathing the cells in the wells of the plate reader (see Methods). Therefore, the values for apparent t_1/2_ of dissociation given in Table 1 for these agonists are an upper limit rather than precise values. However, given that the other agonists examined gave longer apparent t_1/2_ of dissociation values we can conclude that the rate of dissociation from the μ opioid receptor for the agonists tested was fentanyl, alfentanil, DAMGO, morphine > sufentanil > etonitazene = isotonitazene ≥ ohmefentanyl >> carfentanil (Table 1). While in Figure 4d the response to carfentanil was not completely reversed by naloxone 10 μM over the 5 min of naloxone exposure, we subsequently performed a separate series of experiments where naloxone was applied for longer (11 min) and it completely reversed the response to carfentanil (Supplementary Figure 1). In contrast to agonists showing rapid dissociation (e.g. morphine and fentanyl), for agonists exhibiting slow dissociation (e.g. etonitazene and carfentanil) there was a delay from addition of naloxone to the first observable decrease in the amplitude of the agonist response (see inserts in Figure 4a-f and Table 1).

**Figure 4.**
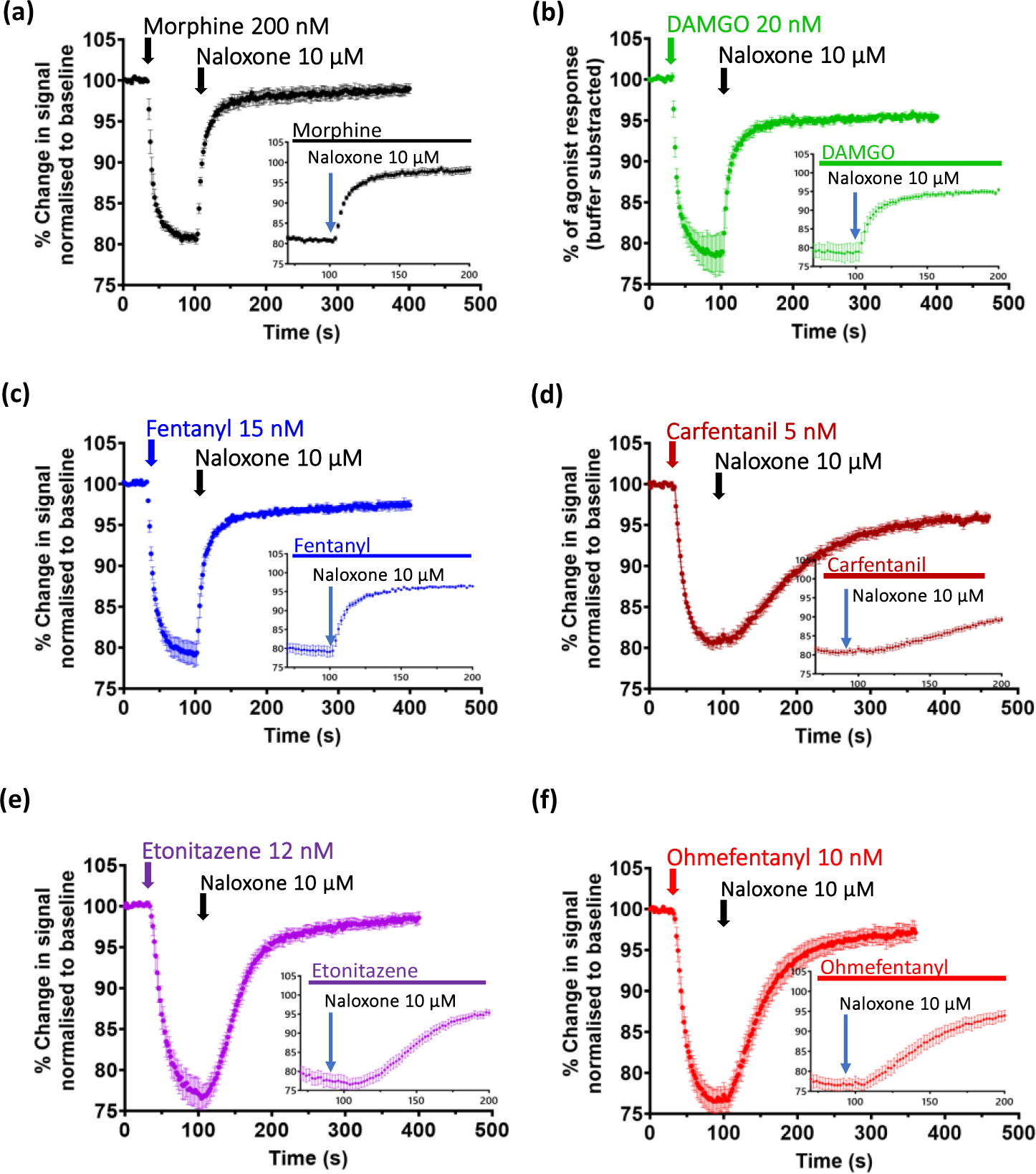
Rate of dissociation of opioid agonists from the μ opioid receptor in AtT20 cells. (**a** – **f**) Pooled experimental data for each opioid agonist showing the change in membrane potential dye fluorescence signal produced by its EC_75_ concentration and subsequent reversal over time by addition of a high concentration of naloxone (10 μM). Traces represent the mean ± SEM of 5 individual experiments for each drug. The inserts show, on an expanded timescale, the responses to naloxone to illustrate the delay to onset of response decay observed with carfentanil, ohmefentanyl and etonitazene but not with morphine, DAMGO and fentanyl. The calculated values for the t_1/2_ of apparent rate of dissociation and the apparent rate constant of dissociation of each are given in Table 1.

Figure 5a shows a graph of the correlation between the apparent rate of agonist dissociation from the receptor and the sensitivity to reversal by naloxone. For the eight agonists studied there was a strong correlation (r = 0.84; p<0.05) between the apparent t_1/2_ of dissociation and reversal by naloxone. Similarly, there was a strong correlation between the apparent t_1/2_ of agonist dissociation and the pA_2_ for antagonism following prior exposure to naloxone (r = 0.92; p<0.05) (Figure 5b). There was a weaker correlation between the t_1/2_ of agonist dissociation from the receptor and agonist potency (r = 0.77; p<0.05) and no observable correlation with lipophilicity (r = 0.56; p>0.05) (Figure 5c & d).

**Figure 5.**
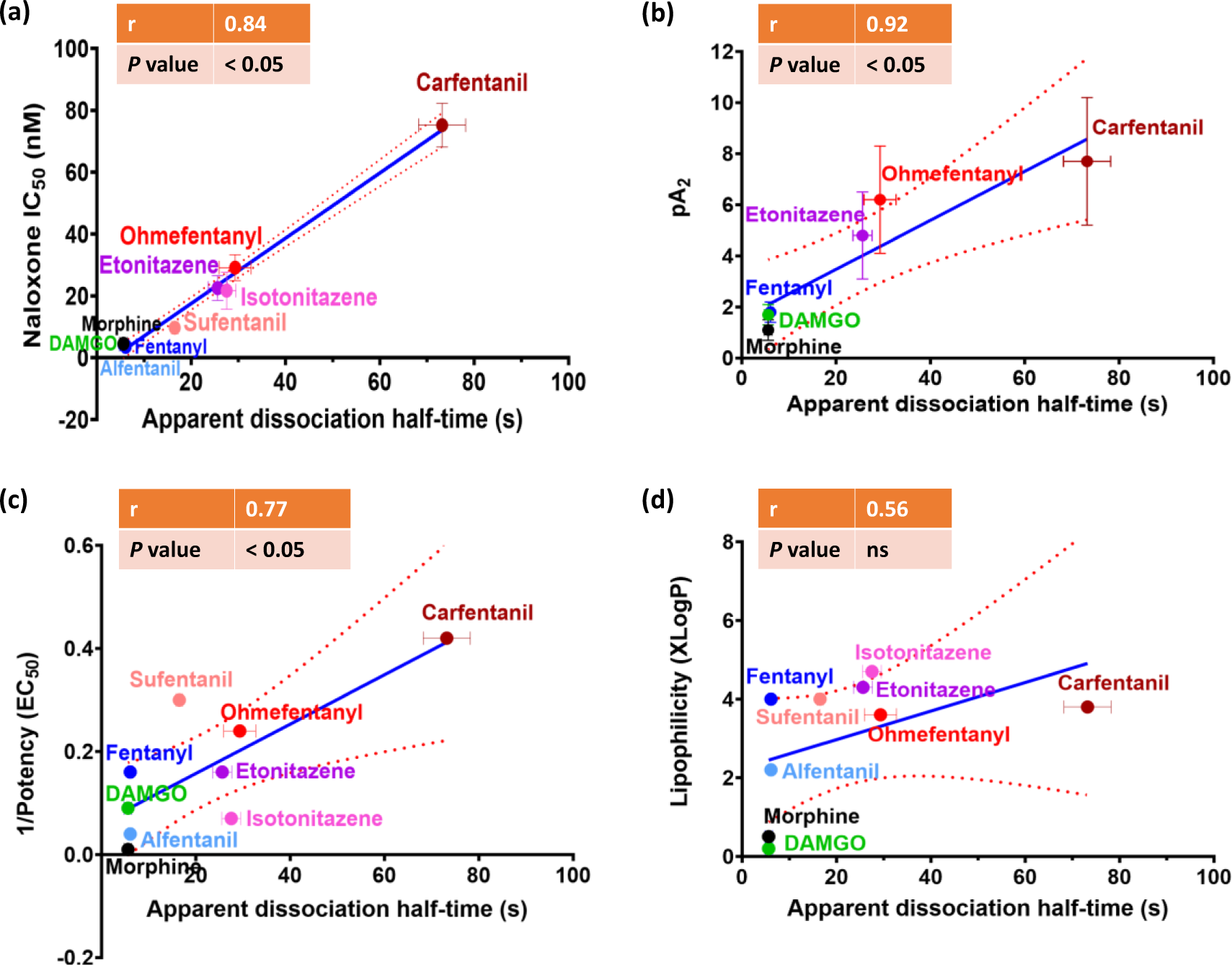
Correlation of apparent agonist dissociation time from the μ opioid receptor with susceptibility to antagonism by naloxone, agonist potency and lipid solubility. The graphs show the correlation between apparent dissociation half time (t_1/2_) with sensitivity to naloxone reversal (**a**), with pA_2_ for naloxone reversal (**b**), with agonist potency (calculated as reciprocal of EC_50_) (**c**) and with agonist lipid solubility (XLogP values obtained from PubChem) (**d**). Data are shown as the mean ± SEM values for each agonist as given in Table 1. The r value and degree of significance obtained by linear regression analysis are indicated on each graph; the dotted lines denote the 95% confidence intervals for the solid regression line.

### 3.4 Opioid agonist signalling bias

As we have previously reported that carfentanil shows bias for β-arrestin translocation over G protein activation (Ramos-Gonzalez, Groom *et al*., 2023) we next sought to examine whether a slow rate of agonist dissociation from the μ opioid receptor was associated with bias for β-arrestin 2 translocation over G protein activation. We compared the ability of DAMGO (reference ligand), morphine, alfentanil, ohmefentanyl, carfentanil, isotonitazene and etonitazene to activate G protein or recruit β-arrestin 2 in HEK293T cells expressing the μ opioid receptor (Figure 6).

**Figure 6.**
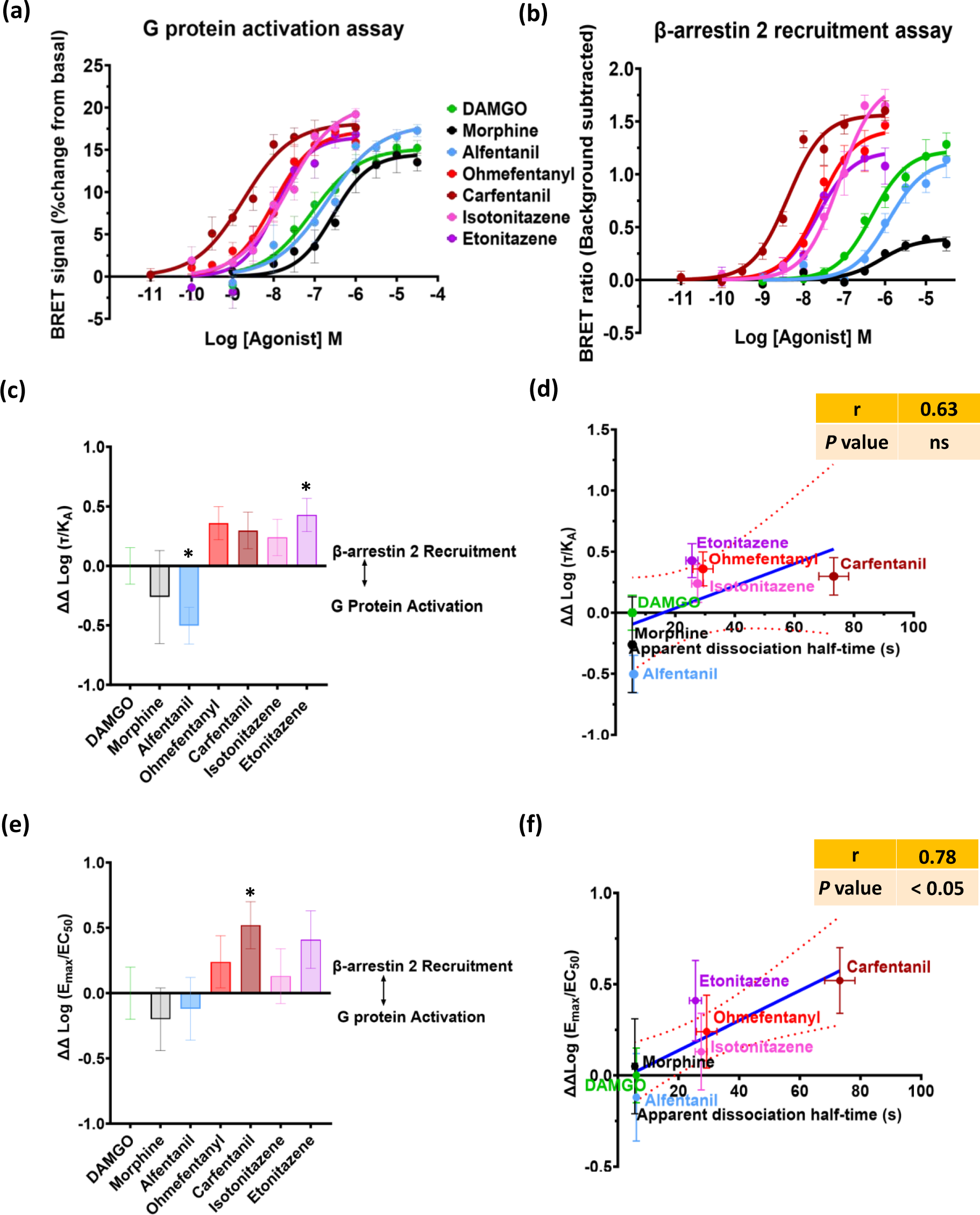
Estimation of opioid agonist bias between G protein activation and β-arrestin 2 translocation at the μ opioid receptor. **a) & b)** Log concentration-response curves for opioid-agonist induced Gi protein activation and β-arrestin 2 translocation in HEK293T cells expressing the μ opioid receptor measured by BRET. Data are expressed as the mean ± SEM, n=5 for each drug; G protein activation is expressed as a % change from basal, β-arrestin 2 recruitment is expressed as raw BRET ratio with the background subtracted. Log concentration-response curves were fitted using nonlinear regression with the Hill slope constrained to 1 and the bottom of the curve constrained to zero. (**c & e**) From the data in **a** & **b** signalling bias was quantified in two ways: using (ΔΔLog(t/K_A_)) and Log(E_max_/EC_50_) (see Methods for details). * denotes statistical significance from 0 (for DAMGO) using a one-sample two-tailed t-test. (**d** & **f**) The graphs show the correlation between apparent dissociation half time (t_1/2_) and (ΔΔLog(t/K_A_)) or ΔΔLog(E_max_/EC_50_) values for the agonists studied. The values shown are the mean ± SEM values for each agonist. The r value and degree of significance obtained by linear regression analysis are indicated on the correlation graphs; the dotted lines denote the 95% confidence intervals for the solid regression line.

The rank order of potency for G protein activation was carfentanil > ohmefentanyl > etonitazene > isotonitazene > DAMGO > alfentanil > morphine (Table 2). Similarly, in the β-arrestin 2 translocation assay, the rank order of potency was carfentanil > ohmefentanyl = etonitazene > isotonitazene > DAMGO > morphine > alfentanil, with morphine exhibiting weak partial agonist activity in this assay (Table 2). Whilst there was some variability in E_max_ values for the agonists in each assay, all three fentanyls and DAMGO signalled with a similar E_max_ for G protein activation (Figure 6a). Isotonitazene, on the other hand, exhibited a significantly higher E_max_ compared to DAMGO. In the β-arrestin 2 translocation assay, the E_max_ values of isotonitazene and carfentanil were significantly higher than that of DAMGO.

**Table 2.** Opioid agonist potency in G_i_ protein activation and β-arrestin 2 translocation BRET assays in HEK293T cells expressing the µ opioid receptor.

|  | G<sub>i</sub> protein assay<sup>†</sup> |  | β-arrestin 2 assay<sup>†</sup> |  |
| --- | --- | --- | --- | --- |
| Agonist | Agonist pEC<sub>50</sub> ± SEM | Agonist E<sub>max</sub> % change from basal | Agonist pEC<sub>50</sub> ± SEM | Agonist E<sub>max</sub> (Raw BRET signal, basal subtracted) |
| Carfentanil | -8.58 ± 0.17* | 17.5 ± 0.96 | -8.38 ± 0.08* | 1.6 ± 0.08* |
| Ohmefentanyl | -8.09 ± 0.12* | 16.6 ± 0.9 | -7.55 ± 0.12* | 1.5 ± 0.07 |
| Etonitazene | -7.90 ± 0.18* | 16.6 ± 1.4 | -7.60 ± 0.04* | 1.3 ± 0.1 |
| Isotonitazene | -7.67 ± 0.18* | 20.5 ± 0.8* | -7.07 ± 0.16* | 1.99 ± 0.2* |
| DAMGO | -6.82 ± 0.06 | 15.5 ± 0.9 | -6.12 ± 0.14 | 1.2 ± 0.1 |
|  | -6.99 ± 0.19 | 14.6 ± 0.6 | -6.33 ± 0.05 | 1.2 ± 0.1 |
| Alfentanil | -6.53 ± 0.22 | 17.5 ± 0.23 | -5.84 ± 0.16 | 1.2 ± 0.1 |
| Morphine | -6.32 ± 0.04 | 14.0 ± 0.7 | -6.17 ± 0.24 | 0.34 ± 0.02* |
|  | -6.38 ± 0.21 | 14.6 ± 1.0 | -6.01 ± 0.12 | 0.39 ± 0.04* |

<sup>†</sup> EC<sub>50</sub>s and maximum responses for each agonist were determined from the concentration-response curves in Figure 6. Values represent the mean ± SEM of 5 different experiments performed in duplicate.
\*Indicates a significant difference (p<0.05) compared to DAMGO using one-way ANOVA with Dunnett's multiple comparisons test. Two sets of experiments were conducted at different times. Each data set was compared to an internal DAMGO control, hence why there are two DAMGO data sets (morphine was also repeated and its two data sets are included for information).

Bias was quantified using both the operational model (Log (τ/K_A_) and Log E_max_/EC_50_. For each method, two statistical tests were performed: one-way ANOVA and t-test (see statistical analysis section). These two statistical tests revealed different outcomes regarding the bias profile of some agonists. For example, carfentanil was β-arrestin 2 biased when bias was quantified using the Log E_max_/EC_50_ method and a one-sample two-tailed t-test. Whereas etonitazene but not carfentanil was β-arrestin 2 biased when bias was quantified using the operational model method and a one-sample two-tailed t-test.

Despite the contrasting statistical findings, correlation analyses of the bias values and apparent dissociation half-times revealed a significant correlation (r = 0.78; p<0.05) between the apparent dissociation half-time of the agonist and ΔΔlog(E_max_/EC_50_) (Figure 6d). In contrast, the correlation was weaker (r = 0.63; p>0.05) and not significant between the apparent dissociation half-time of the agonist and ΔΔLog(t/K_A_) (Figure 6f).

### 3.5 Arrestin binding and agonist off rate

One possibility is that opioid agonist-induced GRK phosphorylation and arrestin binding induces a conformational change in the orthosteric pocket of the μ opioid receptor that decreases agonist dissociation. Such an effect would be most likely to affect the dissociation of those agonists that showed a tendency towards β-arrestin bias. We therefore examined whether the GRK2 & 3 inhibitor, Compound 101 (Cmpd101), which reduces agonist-induced MOR phosphorylation and subsequent β-arrestin binding (Lowe, Sanderson *et al*., 2015) altered agonist dissociation from the μ opioid receptor. DAMGO and carfentanil were examined as the former is a neutral agonist with rapid dissociation and the latter is β-arrestin biased with slow dissociation. Pretreatment of AtT20 cells expressing the μ opioid receptor with Cmpd101 (3 – 30 mM) failed to increase the rate of dissociation of either carfentanil or DAMGO (Figure 7). Indeed, Cmpd101 slightly slowed the rate of dissociation of carfentanil but the effect, while consistent, was not marked and there was no apparent slowing of the rate of dissociation of DAMGO. This suggests that neither GRK phosphorylation and subsequent β-arrestin binding to the μ receptor traps opioid agonists in the orthosteric binding pocket of the receptor.

**Figure 7.**
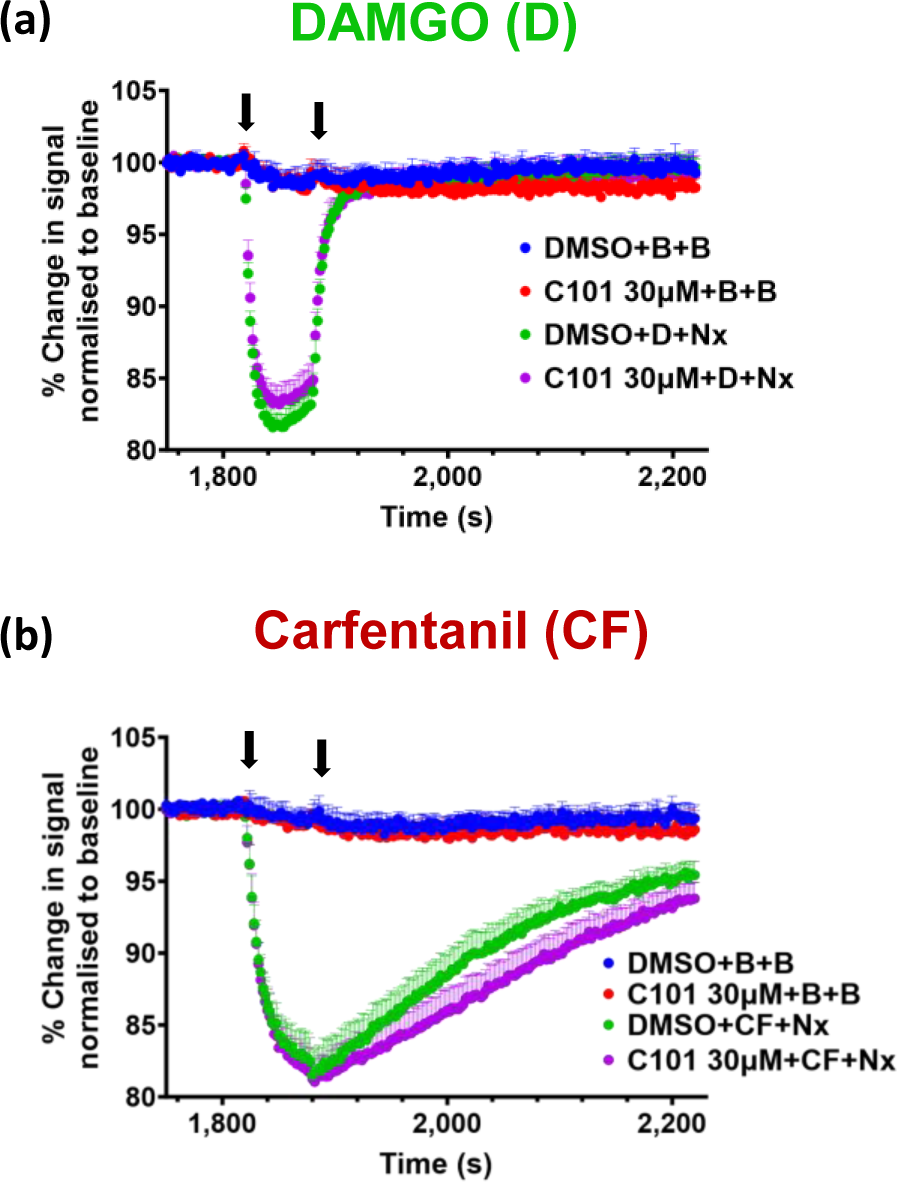
Effect of Compound 101, a G protein receptor kinase (GRK) inhibitor, on the rate of dissociation of agonists from the μ opioid receptor in AtT20 cells. Cells were pretreated with 30 μM Compound 101 (C101) for 30 min before the administration of the EC_75_ concentration of DAMGO (**a**) or carfentanil (**b**). Naloxone (10 µM) was administered 60 s post-agonist. Traces represent the mean ± SEM of 5 individual experiments for each agonist. C101, Compound 101; CF, carfentanil; D, DAMGO; Nx, 10 μM naloxone; B, buffer; DMSO, 0.01%.

## 5. Discussion

When compared to morphine as the standard opioid agonist, the rank order of potency of the fentanyls and nitazenes to produce membrane hyperpolarisation *in vitro* was largely as expected (Tables 1 & 3). These relative potency values agree with those reported previously using a variety of *in vitro* assays involving G protein activation (Emmerson, Clark *et al*., 1996; Toll, Berzetei-Gurske *et al*., 1998; McPherson, Rivero *et al*., 2010; Åstrand, Guerrier *et al*., 2020; Vandeputte, Van Uytfanghe *et al*., 2021; Faouzi, Wang *et al*., 2022; Malcolm, Palkovic *et al*., 2023; Glatfelter, Vandeputte *et al*., 2023; Ramos-Gonzalez, Groom et al., 2023; Vandeputte, Tsai *et al*., 2023).

A key finding of this study was that the rate of drug dissociation from the μ opioid receptor varied significantly between the opioid agonists, alfentanil and fentanyl dissociating rapidly and carfentanil very slowly. The rate of dissociation of carfentanil observed was much faster than that reported by Mann, Samieegohar *et al*. (2022) - (73 s and 2800 s respectively). This may be due to differences in experimental conditions between the studies. Our study was conducted on intact cells in the presence of physiologically relevant concentrations of Na^+^ and guanine nucleotide whereas Mann *et al*. used membrane homogenates and a zero Na^+^ and guanine nucleotide buffer, which would enhance agonist affinity and slow the rate of agonist dissociation (Mohell and Nedergaard, 1985). An early study also measured carfentanil dissociation from the μ opioid receptor in membrane homogenates but in the presence of Na^+^ and guanine nucleotide and reported the off-rate to be 10-fold lower than Mann *et al*. (Titeler, Lyon *et al*., 1989).

The manner in which opioid agonists interact with residues in the orthosteric binding pocket of the µ opioid receptor might explain their different dissociation rates. Whilst a recent cryoEM study (Zhuang, Wang *et al*., 2023) demonstrated that in the orthosteric binding pocket of the µ opioid receptor fentanyl interacts with amino acid residues in transmembrane domains II, III, V, VI and VII, docking experiments in the same study suggested that carfentanil, through its methoxycarbonyl group, forms additional interactions with I298^6.61^, W320^7.35^ and I324^7.39^. We have also observed such interactions following molecular dynamics analysis of carfentanil binding to the active structure of the µ opioid receptor (Ramos-Gonzalez, Groom *et al*., 2023; Ramos-Gonzalez, 2023), whilst lofentanil, which contains a methoxycarbonyl group like carfentanil, also interacts with the above three residues in the orthosteric pocket (Qu, Huang *et al*., 2023). Such additional interactions could decrease the dissociation rate of carfentanil relative to fentanyl and explain carfentanil’s higher affinity for the receptor.

Apart from the orthosteric pocket, it is possible that interactions of carfentanil and similar ligands with other receptor regions, such as a potential vestibule site (Dror, Pan *et al*, 2011), could also hinder the dissociation of carfentanil. The dissociation of LSD from the 5HT_2B_ receptor is greatly slowed by ECL2 of this receptor acting as a “lid” (Wacker, Wang *et al*., 2017), whilst binding to residues in the vestibule region specifically hinders the dissociation of the antagonist tiotropium from the muscarinic M_3_ receptor (Kistemaker, Elzinga *et al*., 2018); further studies will be required to see if such mechanisms operate for carfentanil and other slowly-dissociating ligands from the µ opioid receptor. Potential interactions with the receptor on the way to and from the orthosteric pocket could also enhance the process of agonist rebinding, such that the “on rate” of agonist binding also becomes an important factor contributing to agonist residence time at the receptor (Lane, May *et al*., 2017). We were unable to measure the on rate of agonist binding and receptor activation as the on rate of the response to all the agonists studied was faster than the response time of our assay system. Additionally in conditions of limited diffusion, such as neuronal synapses, the on rate may play a role in prolonging the apparent lifetime of agonist binding through the process of rebinding (Vauquelin and Charlton, 2010).

We have previously reported that carfentanil exhibits bias towards β-arrestin recruitment over G protein activation at the μ opioid receptor (Ramos-Gonzalez, Groom *et al*., 2023) and in the present study we observed that carfentanil and etonitazene exhibited a tendency for β-arrestin bias. However, the GRK inhibitor C101 (Lowe, Sanderson *et al*., 2015) did not enhance the dissociation rate of carfentanil thus excluding the possibility that agonist-induced GRK phosphorylation and subsequent β-arrestin recruitment in some way traps potentially β-arrestin-biased opioid agonists in the orthosteric pocket of the μ opioid receptor slowing their dissociation. The β-arrestin bias of LSD at the 5-HT2B receptor appears to be related to its long residency time in the orthosteric pocket of this receptor (Wacker, Wang *et al*., 2017). Slow agonist dissociation kinetics for dopamine receptor agonists have also been associated with ligand bias (Klein Herenbrink, Sykes *et al*., 2016).

A second important finding of this study was that slowly dissociating agonists such as carfentanil were more resistant to antagonism by naloxone. There have been numerous reports of more naloxone, in the form of multiple or higher doses, being required to reverse overdoses involving fentanyls and nitazenes compared with heroin overdoses (Moe, Godwin *et al*., 2020; Irvine, Oller *et al.,* 2022; Amaducci, Aldy *et al*., 2023). What may be occurring with the fentanyls may in fact be ‘over’ overdose, i.e. far too high a dose of a fentanyl has been unintentionally administered, thus requiring more naloxone for reversal (Rzasa Lynn & Galinkin, 2018). However, in mice higher doses of naloxone were required to reverse non-fatal fentanyl respiratory depression than to reverse morphine (Hill, Santhakumar *et al*., 2020; Elder, Varshneya *et al*., 2023).

In the present *in vitro* study higher concentrations of naloxone were required to antagonise some fentanyls and nitazenes irrespective of whether the naloxone was applied before or after the agonist. Surprisingly, given our *in vivo* mouse respiration data (Hill, Santhakumar *et al*., 2020) fentanyl showed similar reversal by naloxone as morphine *in vitro*. For all of the agonists studied antagonism was indeed surmountable i.e. there was a parallel shift to the right of the concentration-response curve in the presence of increasing concentrations of naloxone with no decrease in maximum agonist response. However, the slopes of the Schild plots for the fentanyls and nitazenes were lower than unity and the intercepts on the X axis (pA2 values) increased in parallel with the reduction in slope. The results of the Schild analysis are incompatible with competitive agonist-antagonist interaction at the orthosteric binding site of a single receptor type. Given that the AtT20 cells we used in our assay only expressed μ opioid receptors we therefore conclude that naloxone antagonism of fentanyls and nitazenes at the μ opioid receptor is pseudo-competitive in nature. The strong correlation between the rate of agonist dissociation from the receptor and susceptibility to naloxone antagonism indicates that slow agonist dissociation impairs naloxone antagonism, rendering it pseudo-competitive. In a similar way the *in vivo* actions of the slowly dissociating CB_1_ cannabinoid agonist HU-210 are less susceptible to antagonism by rimonabant than other faster-dissociating agonists at the CB_1_ receptor (Hruba and McMahon, 2014). Other potential mechanisms that might contribute to reduced sensitivity to naloxone include differential binding of some opioid agonists to orthosteric and vestibule sites (discussed above) on the receptor, the latter reducing access of the antagonist to the orthosteric site. Also, highly lipophilic agonists may be able to access the orthosteric pocket of the receptor both by the aqueous route and through the transmembrane helices of the receptor while naloxone uses only the aqueous route (Kelly, Sutcliffe *et al*., 2023; Sutcliffe, Corey *et al*., 2022). These potential mechanisms would result in nonequilibrium conditions and reduce antagonist sensitivity (Kenakin, 1982).

Across a number of *in vivo* studies of antinociception the relative potency of fentanyls and nitazenes compared to morphine is much higher than that observed using *in vitro* assays (Table 3). Furthermore, the change in relative potency between *in vitro* and *in vivo* assays was not consistent across the range of fentanyls tested. For fentanyl the difference was around 20-fold but for carfentanil it was 350-fold. Fentanyl and nitazene analogues are highly lipophilic, but this cannot fully explain the disparity between their *in vitro* and *in vivo* relative potencies as carfentanil and fentanyl are of similar lipophilicity (XlogP values of 4.0 and 3.8 respectively), but the enhanced relative potency of carfentanil *in vivo* was much greater. Fentanyl, alfentanil and morphine are substrates for P-glycoprotein-mediated extrusion from the brain (Dagenais, Graff *et al*., 2004; Kalvas, Olson *et al*., 2007; Yu, Yuan *et al*., 2018; Martins, Loos *et al*., 2023). However it seems unlikely that the *in vitro-in vivo* relative potency disparity between fentanyl/alfentanil and carfentanil/ohmefentanyl/isotonitazene/etonitazene could be attributed to P-glycoprotein excluding fentanyl and alfentanil from the brain more effectively, as pharmacological blockade or genetic deletion of P-glycoprotein only increases the brain levels of fentanyl and alfentanil by 3-fold (Kalvas, Olson *et al*., 2007; Yu, Yuan *et al*., 2018). The mechanisms responsible for the enhanced potency of fentanyls and nitazenes *in vivo*, as well as the much greater *in vivo* potency of carfentanil relative to fentanyl remain unexplained.

**Table 3.**
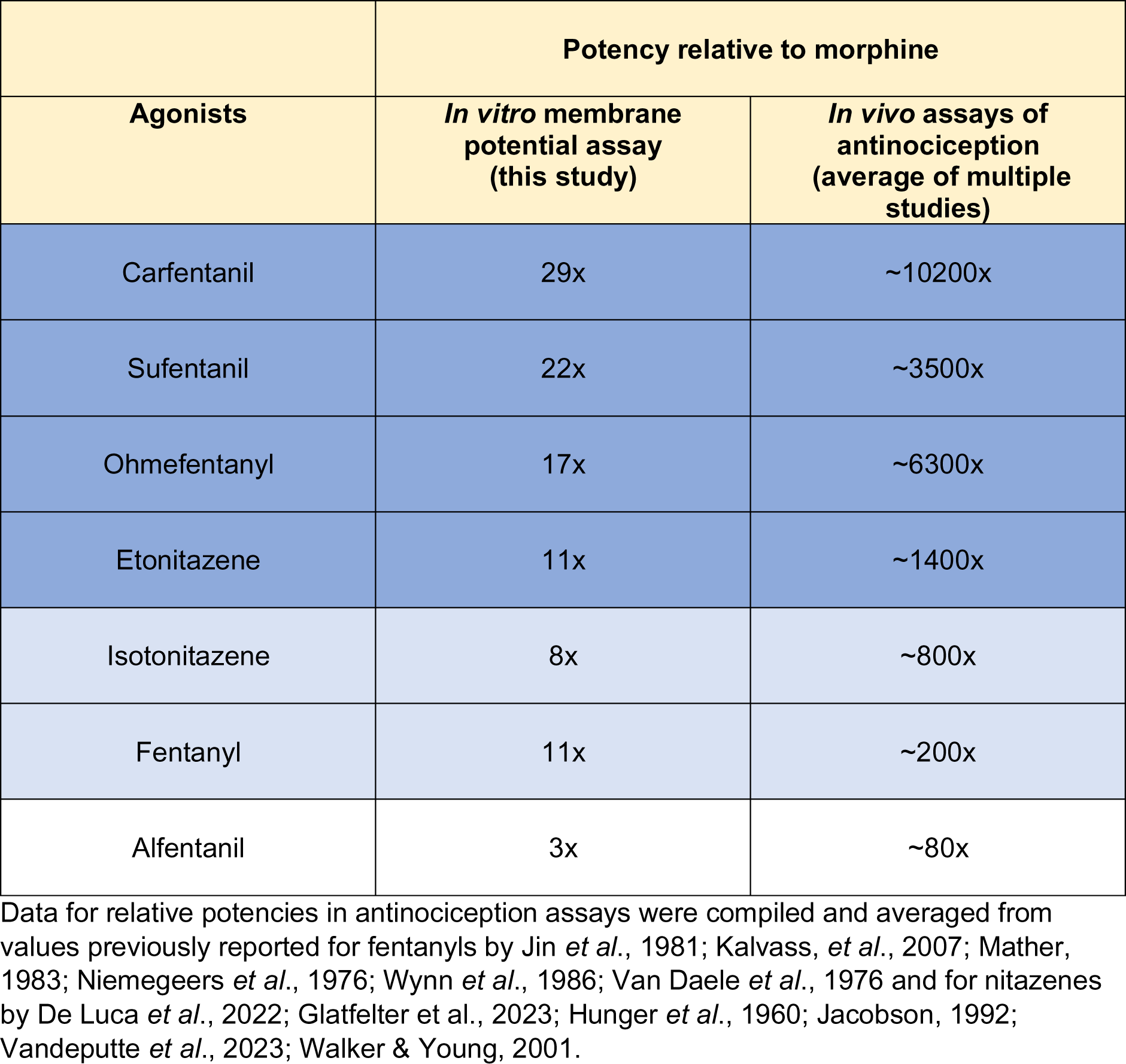
Comparison of relative potencies of fentanyls and nitazenes to morphine in an *in vitro* assay and *in vivo* antinociception assays.

## 5. Conclusions

Our data confirm the view that in overdoses involving fentanyls and nitazenes higher doses of the antidote naloxone may be required for reversal than those normally used to reverse heroin overdose. With slowly dissociating μ opioid receptor agonists antagonism by naloxone becomes pseudo competitive. This indicates that the kinetics of the agonist should be factored in when evaluating the nature of antagonism.

## Acknowledgments

This research was not preregistered with an analysis plan in an independent, institutional registry.

## Abbreviations

5-HT: 5-hydroxy tryptamine
ANOVA: analysis of variance
BRET: Bioluminescence Resonance Energy Transfer
Cmpd101: Compound 101
Cryo-EM: cryogenic electron microscopy
DAMGO: D-Ala^2^, *N*-MePhe^4^, Gly-ol]-enkephalin
DMEM: Dulbecco’s modified Eagle’s medium
ECL: extracellular loop
FBS: fetal bovine serum
GIRK: G protein-coupled inwardly rectifying potassium channel
GPCR: G protein-coupled receptor
GRK: G protein-coupled receptor kinase
HEK293T: human embryonic kidney 293
IUPHAR: International Union of Basic and Clinical Pharmacology
LSD: lysergic acid diethylamide
RlucII: *Renilla* luciferase II
S.D.: standard deviation
S.E.M: standard error of the mean.

## DATA AVAILABILITY STATEMENT

The data that support the findings of this study are available from the corresponding author upon reasonable request.

## DECLARATION OF TRANSPARENCY AND SCIENTIFIC RIGOUR

This Declaration acknowledges that this paper adheres to the principles for transparent reporting and scientific rigour of preclinical research as stated in the BJP guidelines for Design and Analysis and Animal Experimentation, and as recommended by funding agencies, publishers and other organisations engaged with supporting research.

## Author Contributions

*Participated in research design*: Norah Alhosan, Damiana Cavallo, Eamonn Kelly and Graeme Henderson. *Supplied AtT20FlpInMOR cells, advised on membrane potential assay procedure*: Maria Santiago. *Conducted experiments*: Norah Alhosan, Damiana Cavallo. *Performed data analysis*: Norah Alhosan, Eamonn Kelly and Graeme Henderson. *Contributed to the writing of the manuscript*: all authors.

## Supplementary figure

**Supplementary Figure 1.**
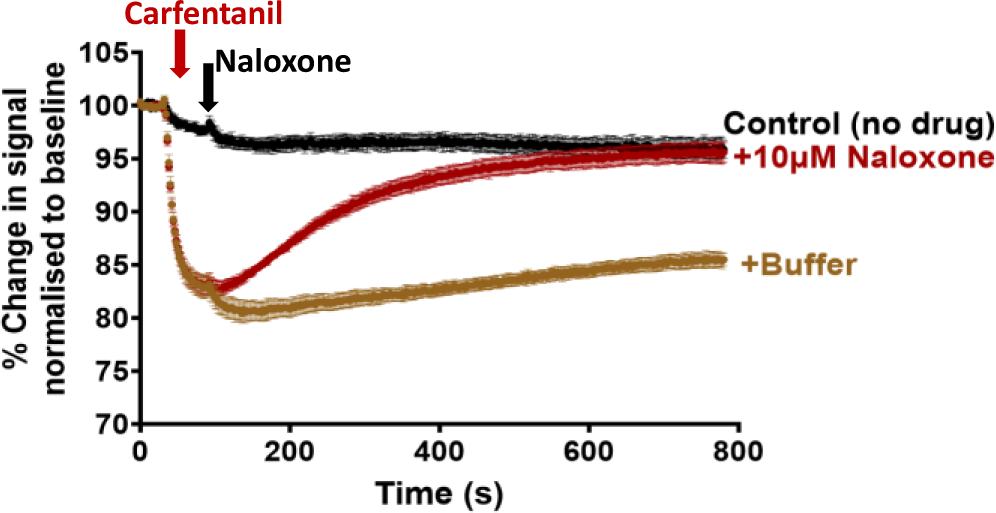
Complete reversal by naloxone of the carfentanil-induced hyperpolarisation of μ opioid receptor-expressing AtT20 cells. Pooled experimental data showing the change in membrane potential dye fluorescence signal produced by the EC_75_ concentration of carfentanil and subsequent reversal over time by addition of a high concentration of naloxone (10 μM). Traces represent the mean ± SEM of 5 individual experiments.

## References

Alexander, S. P., Christopoulos, A., Davenport, A. P., Kelly, E., Mathie, A., Peters, J. A., … Ye, R. D. (2021). The Concise Guide to Pharmacology 2021/22: G protein-coupled receptors. British Journal of Pharmacology, 178(S1), S27–S156. 10.1111/bph.15538.

Amaducci, A., Aldy , K., Campleman, S.L., Li, S., Meyn, A., Abston, S., … Manini, A.F. (2023). Naloxone use in novel potent opioid and fentanyl overdoses in emergency department patients. Toxicology Investigators Consortium Fentalog Study Group. JAMA Network Open. 6(8):e2331264. 10.1001/jamanetworkopen.2023.31264.

Åstrand, A., Guerrieri, D., Vikingsson, S., Kronstrand, R., & Green, H. (2020). *In vitro* characterization of new psychoactive substances at the μ-opioid, CB1, 5HT1A, and 5-HT2A receptors—On-target receptor potency and efficacy, and off-target effects. Forensic Science International, 317, 110553. 10.1016/j.forsciint.2020.110553.

Black, J.W. & Leff, P. (1983). Operational models of pharmacological agonism. Proceedings Royal Society Lond B, 220:141–62. 10.1098/rspb.1983.0093

Caulkins, J.P., Tallaksen, A., Taylor, J., Kilmer, B., & Reuter P. (2024). The Baltic and Nordic responses to the first Taliban poppy ban: Implications for Europe & synthetic opioids today. International Journal of Drug Policy 124, 104314. 10.1016/j.drugpo.2023.104314.

Centers for Disease Control and Prevention (CDC). (2022). Understanding the Opioid Overdose Epidemic. Retrieved from https://www.cdc.gov/opioids/basics/epidemic.html

Corbett, A. D., Henderson, G., McKnight, A. T., & Paterson, S. J. (2006). 75 years of opioid research: the exciting but vain quest for the Holy Grail. British Journal of Pharmacology, 147 Suppl(Suppl 1), S153–62. 10.1038/sj.bjp.0706435

Curtis, M. J., Alexander, S., Cirino, G., Docherty, J. R., George, C. H., Giembycz, M. A., … Ahluwalia, A. (2018). Experimental design and analysis and their reporting II: Updated and simplified guidance for authors and peer reviewers. British Journal of Pharmacology, 175, 987–993. 10.1111/bph.14153

Curtis, M. J., Alexander, S. P. H., Cirino, G., George, C. H., Kendall, D. A., Insel, P. A., … Ahluwalia, A. (2022). Planning experiments: Updated guidance on experimental design and analysis and their reporting III. In British Journal of Pharmacology, 179, 3907–3913. 10.1111/bph.15868

Dagenais, C., Graff, C. L., & Pollack, G. M. (2004). Variable modulation of opioid brain uptake by P-glycoprotein in mice. Biochemical Pharmacology, 67(2), 269–276. 10.1016/j.bcp.2003.08.027.

De Luca, M. A., Tocco, G., Mostallino, R., Laus, A., Caria, F., Musa, A., Pintori, N., Ucha, M., Poza, C., Ambrosio, E., Di Chiara, G., & Castelli, M. P. (2022). Pharmacological characterization of novel synthetic opioids: Isotonitazene, metonitazene, and piperidylthiambutene as potent μ opioid receptor agonists. Neuropharmacology, 221, 109263. 10.1016/j.neuropharm.2022.109263

Dror, R. O., Pan, A. C., Arlow, D. H., Borhani, D. W., Maragakis, P., Shan, Y., Xu, H., & Shaw, D. E. (2011). Pathway and mechanism of drug binding to G protein-coupled receptors. Proceedings of the National Academy of Sciences, 108(32), 13118–13123. 10.1073/pnas.1104614108

Elder, H. J., Varshneya, N. B., Walentiny, D. M., & Beardsley, P. M. (2023). Amphetamines modulate fentanyl-depressed respiration in a bidirectional manner. Drug and Alcohol Dependence, 243, 109740. 10.1016/J.DRUGALCDEP.2022.109740

Emmerson, P. J., Clark, M. J., Mansour, A., Akil, H., Woods, J. H., & Medzihradsky, F. (1996). Characterization of opioid agonist efficacy in a C6 glioma cell line expressing the μ opioid receptor. Journal of Pharmacology and Experimental Therapeutics, 278, :1121–1127.

Faouzi, A., Wang, H., Zaidi, S. A., DiBerto, J. F., Che, T., Qu, Q., Robertson, M. J., Madasu, M. K., El Daibani, A., Varga, B. R., Zhang, T., Ruiz, C., Liu, S., Xu, J., Appourchaux, K., Slocum, S. T., Eans, S. O., Cameron, M. D., Al-Hasani, R., … Majumdar, S. (2022). Structure-based design of bitopic ligands for the µ-opioid receptor. Nature 2022 613:7945, 613(7945), 767–774. 10.1038/s41586-022-05588-y

Giraudon, I., Abel-Ollo, Ki, Vanaga-Arāja, D., Heudtlass, P., & Griffiths, P. (2024). Nitazenes represent a growing threat to public health in Europe. Lancet Public Health 9, 4e216. 10.1016/S2468-2667(24)00024-0

Glatfelter, G.C., Vandeputte, M.M., Chen, L., Walther, D., Tsai, M.M., Shi, L., Stove, C.P., & Baumann, M.H. (2023). Alkoxy chain length governs the potency of 2-benzylbenzimidazole ‘nitazene’ opioids associated with human overdose. Psychopharmacology (Berl). 240,2573–2584. 10.1007/s00213-023-06451-2.

Hill, R., Disney, A., Conibear, A., Sutcliffe, K., Dewey, W., Husbands, S., … & Henderson G. (2018). The novel μ-opioid receptor agonist PZM21 depresses respiration and induces tolerance to antinociception. British Journal of Pharmacology, 175, 2653–2661. 10.1111/bph.14224.

Hill, R., Santhakumar, R., Dewey, W., Kelly, E., & Henderson G. (2020). Fentanyl depression of respiration: Comparison with heroin and morphine. British Journal of Pharmacology, 177, 254–266. 10.1111/bph.14860.

Holland, A., Copeland, C.S., Shorter, G.W., Conolly, D.J., Wiseman, A., Mooney, J., Fenton, K., & Harris, M. (2024). Nitazenes—heralding a second wave for the UK drug-related death crisis? Lancet Public Health, 9, E71–72. 10.1016/S2468-2667(24)00001-X

Hunger, A., Kebrle, J., Rossi, A., & Hoffmann, K. (1960). Benzimidazol-derivate und verwandte heterocyclen III. Synthese von 1-aminoalkyl-2-benzyl-nitro-benzimidazolen. Helvetica Chimica Acta, 43, 1032–1046. 10.1002/hlca.19600430412.

Irvine, M.A., Oller, D., Boggis, J., Bishop, B., Coombs, D., Wheeler, E., … Green, T.C. (2022) Estimating naloxone need in the USA across fentanyl, heroin, and prescription opioid epidemics: a modelling study. Lancet Public Health, 7(3):e210–e218. 10.1016/S2468-2667(21)00304-2.

Jacobson, A. E. (1992). Biological evaluation of compounds for their physical dependence potential and abuse liability, XIV. Animal Testing Committee of the Committee on Problems of Drug Dependence, Inc. (1991). NIDA Research Monograph, 119, 490–512.

Jin, W. Q., Xu, H., Zhu, Y. C., Fang, S. N., Xia, X. L., Huang, Z. M., Ge, B. L., & Chi, Z. 296 Q. (1981). Studies on synthesis and relationship between analgesic activity and opioid receptor affinity for 3-methyl fentanyl derivatives. Scientia Sinica, 24(5), 710–720. http://www.ncbi.nlm.nih.gov/pubmed/6264594.

Kalvass, J. C., Olson, E. R., Cassidy, M. P., Selley, D. E., & Pollack, G. M. (2007). Pharmacokinetics and pharmacodynamics of seven opioids in P-glycoprotein-competent mice: Assessment of unbound brain EC50, correlation of in vitro, preclinical, and clinical data. Journal of Pharmacology and Experimental Therapeutics, 323, 346–355. 10.1124/JPET.107.119560.

Kelly, E., Sutcliffe, K., Cavallo, D., Ramos-Gonzalez, N., Alhosan, N., & Henderson, G. (2023). The anomalous pharmacology of fentanyl. British Journal of Pharmacology, 180, 797–812. 10.1111/BPH.15573.

Kenakin, T. P. (1982). The Schild regression in the process of receptor classification. Canadian Journal of Physiology and Pharmacology, 60, 249–265. 10.1139/y82-036.

Kistemaker, L. E. M., Elzinga, C. R. S., Tautermann, C. S., Pieper, M. P., Seeliger, D., Alikhil, S., …Gosens, R. (2019). Second M_3_ muscarinic receptor binding site contributes to bronchoprotection by tiotropium. British Journal of Pharmacology, 176(16), 2864–2876. 10.1111/BPH.14707

Klein Herenbrink, C., Sykes, D.A., Donthamsetti, P., Canals, M., Coudrat, T. Shonberg, J., …& Lane, J.R. (2016). The role of kinetic context in apparent biased agonism at GPCRs. Nature Communications, 24:7:10842. 10.1038/ncomms10842.

Knapman, A., Santiago, M., & Connor, M. (2014). Buprenorphine signalling is compromised at the N40D polymorphism of the human μ opioid receptor in vitro. British Journal of Pharmacology, 171(18), 4273. 10.1111/BPH.12785

Kolb, P., Kenakin, T., Alexander, S. P. H., Bermudez, M., Bohn, L. M., Breinholt, C. S., … & Gloriam, D. E. (2022). Community guidelines for GPCR ligand bias: IUPHAR review 32. British Journal of Pharmacology, 179(,3651–3674. 10.1111/BPH.15811

Lane, J.R. May, L.T., Parton, R.G., Sexton, P.M., & Christopoulos, A. (2017) A kinetic view of GPCR allostery and biased agonism. Nature Chemical Biology. 13:929–937. 10.1038/nchembio.2431.

Lowe, J. D., Sanderson, H. S., Cooke, A. E., Ostovar, M., Tsisanova, E., Withey, S. L, … & Bailey, C. P. (2015). Role of G protein-coupled receptor kinases 2 and 3 in μ-opioid receptor desensitization and internalization. Molecular Pharmacology, 88(2), 347–356. 10.1124/MOL.115.098293/-/DC1

McPherson, J., Rivero, G., Baptist, M., Llorente, J., Al-Sabah, S., Krasel, C., … & Kelly, E. (2010). μ-Opioid receptors: correlation of Agonist efficacy for signalling with ability to activate internalization. Molecular Pharmacology, 78(4), 756–766. 10.1124/MOL.110.066613

Malcolm, N. J., Palkovic, B., Sprague, D. J., Calkins, M. M., Lanham, J. K., Halberstadt, A. L., Stucke, A. G., & McCorvy, J. D. (2023). μ-Opioid receptor selective superagonists produce prolonged respiratory depression. IScience, 26(7), 107121. 10.1016/J.ISCI.2023.107121

Mann, J., Samieegohar, M., Chaturbedi, A., Zirkle, J., Han, X., Ahmadi, S. F., … & Li, Z. (2022). Development of a translational model to assess the impact of opioid overdose and naloxone dosing on respiratory depression and cardiac arrest. Clinical Pharmacology and Therapeutics, 112, 1020–1032. 10.1002/CPT.2696

Martins, M.L.F., Loos, N.H.C., El Yattouti, M. Offeringa, L., Heydari, P., Hillebrand, M.J.X., … & Schinkel, A.H. (2023). P-glycoprotein (MDR1/ABCB1) restricts brain penetration of the main active heroin metabolites 6-monoacetylmorphine (6-MAM) and morphine in mice. Pharmacology Research. 40:1885–1899. 10.1007/s11095-023-03545-6.

Mather, L. E. (1983). Clinical pharmacokinetics of fentanyl and its newer derivatives. Clinical Pharmacokinetics, 8, 422–446. 10.2165/00003088-198308050-00004

Moe, J., Godwin, J., Purssell, R., O’sullivan, F., Hau, J. P., Purssell, E., Curran, J., Doyle-Waters, M. M., Brasher, P. M. A., Buxton, J. A., & Hohl, C. M. (2020). Naloxone dosing in the era of ultra-potent opioid overdoses: A systematic review. Canadian Journal of Emergency Medicine, 22(2), 178–186. 10.1017/cem.2019.471

Mohell, N. & Nedergaard, J. (1985) Effects of guanine nucleotides and cations on agonist affinity of alpha 1-adrenoceptors in brown adipose tissue. European Journal of Pharmacology, 115:231–240. 10.1016/0014-2999(85)90695-8

Niemegeers, C. J., Schellekens, K. H., Van Bever, W. F., & Janssen, P. A. (1976). Sufentanil, a very potent and extremely safe intravenous morphine-like compound in mice, rats and dogs. Arzneimittel-Forschung, 26, 1551–1556. PMID: 12772

Qu, Q., Huang, W., Aydin, D., Paggi, J. M., Seven, A. B., Wang, H., … & Skiniotis, G. (2022). Insights into distinct signaling profiles of the µOR activated by diverse agonists. Nature Chemical Biology, 19, 423–430. 10.1038/s41589-022-01208-y

Ramos-Gonzalez, N. (2023). An investigation of the activation of µ opioid receptors by fentanyls using in silico and in vitro approaches — University of Bristol. https://research-information.bris.ac.uk/en/studentTheses/an-investigationof-the-activation-of-µ-opioid-receptors-by-fenta

Ramos-Gonzalez, N., Groom, S., Sutcliffe, K. J., Bancroft, S., Bailey, C. P., Sessions, R. B., Henderson, G., & Kelly, E. (2023). Carfentanil is a β-arrestin-biased agonist at the μ opioid receptor. British Journal of Pharmacology, 180, 2341–2360. 10.1111/BPH.16084

Ritter, J., Flower, R. J., Henderson, G., Loke, Y. K., MacEwan, D. J., Robinson, E. S. J., & Fullerton, J. (2024). Rang and Dale’s ‘Pharmacology’ 10^th^ edition (2024). How drugs act: general principles. Elsevier (London). ISBN: 9780323873956.

Rzasa Lynn, R., & Galinkin, J. L. (2018). Naloxone dosage for opioid reversal: current evidence and clinical implications. Therapeutic Advances in Drug Safety, 9, 63–88. 10.1177/2042098617744161.

Sutcliffe, K. J., Corey, R. A., Alhosan, N., Cavallo, D., Groom, S., Santiago, M., … & Kelly, E. (2022). Interaction with the lipid membrane influences fentanyl pharmacology. Advances in Drug and Alcohol Research, 2. 10.3389/ADAR.2022.10280.

Suzuki, J., & El-Haddad, S. (2017). A review: Fentanyl and non-pharmaceutical fentanyls. Drug and Alcohol Dependence, 171, 107–116. 10.1016/J.DRUGALCDEP.2016.11.033.

Titeler, M., Lyon, R. A., Kuhar, M. J., Frost, J. F., Dannals, R. F., Leonhardt, S., … & Struble, R. G. (1989). μ Opiate receptors are selectively labelled by [^3^H]carfentanil in human and rat brain. European Journal of Pharmacology, 167, 221–228. 326. 10.1016/0014-2999(89)90582-7.

Toll, L., Berzetei-Gurske, I. P., Polgar, W. E., Brandt, S. R., Adapa, I. D., Rodriguez, L., … & Auh, J. S. (1998). Standard binding and functional assays related to medications development division testing for potential cocaine and opiate narcotic treatment medications. NIDA Research Monograph, 178, 440–466. PMID: 9686407.

Van Daele, P. G., De Bruyn, M. F., Boey, J. M., Sanczuk, S., Agten, J. T., & Janssen, P. A. (1976). Synthetic analgesics: N-(1-[2-arylethyl]-4-substituted 4-piperidinyl) N-arylalkanamides. Arzneimittel-Forschung, 26, 1521–1531. 10.1002/chin.197646236

Vandeputte, M. M., Van Uytfanghe, K., Layle, N. K., St. Germaine, D. M., Iula, D. M., & Stove, C. P. (2021). Synthesis, chemical characterization, and μ-opioid receptor activity assessment of the emerging group of “nitazene” 2-benzylbenzimidazole synthetic opioids. ACS Chemical Neuroscience, 12, 1241–1251. https://pubs.acs.org/doi/full/10.1021/acschemneuro.1c00064.

Vandeputte, M. M., Tsai, M.-H. M., Chen, L., Glatfelter, G. C., Walther, D., Stove, C. P., Shi, L., & Baumann, M. H. (2023). Comparative neuropharmacology of structurally distinct non-fentanyl opioids that are appearing on recreational drug markets worldwide. Drug and Alcohol Dependence, 249, 109939. 10.1016/J.DRUGALCDEP.2023.109939.

Vauquelin, G., & Charlton, S. J. (2010). Long-lasting target binding and rebinding as mechanisms to prolong *in vivo* drug action. British Journal of Pharmacology, 161, 488–508. 10.1111/j.1476-5381.2010.00936.x

Wacker, D., Wang, S., McCorvy, J. D., Betz, R. M., Venkatakrishnan, A. J., Levit, A., … & Roth, B. L. (2017). Crystal structure of an LSD-bound human serotonin receptor. Cell, 168, 377–389.e12. 10.1016/j.cell.2016.12.033.

Walker, E.A., Young, A.M. (2001). Differential tolerance to antinociceptive effects of mu opioids during repeated treatment with etonitazene, morphine, or buprenorphine in rats. Psychopharmacology (Berl). 154:131–42. 10.1007/s002130000620.

Wynn, R. L., Ford, R. D., Mccourt, P. J., Ramkumar, V., Bergman, S. A., & Rudo, F. G. (1986). Rabbit tooth pulp compared to 55°C mouse hot plate assay for detection of antinociceptive activity of opiate and nonopiate central analgesics. Drug Development Research, 9, 233–239. 10.1002/ddr.430090308.

Yu, C., Yuan, M., Yang, H., Zhuang, X., & Li, H. (2018). P-glycoprotein on blood-brain barrier plays a vital role in fentanyl brain exposure and respiratory toxicity in rats. Toxicological Sciences, 164, 353–362. 10.1093/toxsci/kfy093.

Zhuang, Y., Wang, Y., He, B., He, X., Zhou, X. E., Guo, S., … Xu, H. E. (2022). Molecular recognition of morphine and fentanyl by the human μ-opioid receptor. Cell, 185, 4361–4375.e19. 10.1016/J.CELL.2022.09.041.

